# Predicting the Efficacy of Myopia Control Spectacle Lenses by Simulating Their Optical Effect on the Retina

**DOI:** 10.64898/2026.10.06.756491

**Authors:** Samy Hamlaoui, Sarah Goethals, Konogan Baranton, Thomas Bessenay, Sylvain Mercier, Olivier Marre

## Abstract

Slowing myopia progression is central to addressing a growing public health concern. Myopia control spectacle lenses with diverse optical designs have been proposed for this purpose, and show varying degrees of clinical efficacy. Relating their design to their efficacy remains an open challenge. Here we show that it is possible to predict their clinical efficacy from their optical design, by reconstructing their impact on the retinal image. We simulated how several lens designs affect the retinal image using a combined lens and eye optical model. By assuming that the retina extracts specific features from the retinal image to modulate eye growth, we could predict the cumulative absolute reduction in axial elongation (CARE) of lens designs with very different rationales and microstructures. Our results show that different myopia control spectacle lenses can be compared on a common optical basis, and that their efficacy is predictable from their optical properties. This work paves the way for rational design of next-generation lenses and for adapting designs to each patient’s ocular optics.

## 1. Introduction

It has been predicted that half of the worldwide population will be myopic by 2050, and that a substantial fraction will have high myopia (Holden et al., 2016). High myopia is associated with an increased risk of several potentially sight-threatening ocular complications, including myopic macular degeneration, retinal detachment, glaucoma, and cataract (Haarman et al., 2020). Slowing axial elongation, and thereby myopia progression, is therefore an important response to an impending public health concern.

Eye growth is modulated by visual input. The eye responds differently to myopic and hyperopic defocus. Myopic defocus places the focal plane in front of the retina and can slow eye growth. This regulation operates locally within the retina (Smith et al., 2009). One strategy to slow myopia progression is to alter the visual input received by the patient. Several myopia control (MC) spectacle lenses have been developed for this purpose, with encouraging but heterogeneous results (Schmidt et al., 2025). Early MC spectacle lenses were designed to create myopic defocus in the periphery (Sankaridurg et al., 2010). However, other optical strategies may also slow eye growth. Neitz & Neitz (2024) proposed reducing peripheral contrast with light-scattering elements, and Su et al. (2024) showed that lenslets focusing both in front of and behind the retina can both reduce myopia progression. Optical characterization studies have examined these new lenses (Gantes-Nuñez et al., 2023; Jaskulski et al., 2020), and alternative mechanistic explanations have also been proposed (Gawne et al., 2025; Guggenheim & Terry, 2025). Together, these strategies point to two broad and probably complementary descriptions of the optical signal delivered by MC spectacle lenses: one based on the sign and magnitude of defocus, the other on retinal contrast reduction, which may lower retinal activity and has been proposed as a pathway for reduced eye growth (Neitz & Neitz, 2025). These descriptions are not mutually exclusive, since a lenslet array that creates peripheral defocus also redistributes light and therefore changes retinal-image contrast (Papadogiannis et al., 2023).

Spectacle lenses with very different optical designs can thus slow myopia progression, but with heterogeneous efficacies. Because these designs rely on different optical principles, there is no common basis on which to compare them or relate them to clinical outcomes.

Because the retina is the first stage that processes visual input to determine defocus sign and modulate eye growth, the efficacy of MC spectacle lenses depends largely on the image formed at the photoreceptor layer. However, the mechanisms by which an optical signal is transformed into a biological signal remain unclear, and there are still several hypotheses about the features that are used by the retina to determine the sign of defocus (Schaeffel & Swiatczak, 2024).

Here we reconstructed the retinal image formed by several MC spectacle lenses, using a combination of optical measurements and modeling. This characterization provides a common basis for comparing lens designs with different optical strategies. We showed that the different lens designs produced distinct point spread functions (PSFs) at the retina. From the simulated retinal images, we then extracted local spatial contrast (LSC), a feature that the retina could plausibly use to determine the sign of defocus and regulate eye growth, and found that it was strongly associated with reported clinical efficacy. We further examined how this relationship varied with spatial integration scale and retinal eccentricity under two representative gaze configurations. These results show that MC spectacle lenses can be compared on a common optical basis, and that retinal image metrics may support evidence-based lens design.

## 2. Results

We first developed a framework to simulate the images of indoor scenes formed on the retina through MC spectacle lenses and the eye optics. We then extracted image-based metrics from these simulated retinal images and assessed whether they were associated with reported clinical efficacy. We focused the analysis on MC spectacle lenses that do not substantially alter the accommodative response, unlike progressive and bifocal MC spectacle lenses (see methods).

### 2.1 Simulating the images received by the retina

To simulate retinal PSFs for each MC spectacle lens-eye system, we first measured each MC surface or reconstructed it from published data. For each lens, we then assembled the lens-eye system and computed a polychromatic through-focus set of retinal PSFs, which we convolved with indoor scenes to obtain retinal images. We calculated LSC, which quantifies how much contrast varies within a small retinal region, for each retinal image and compared its spatial distribution with that obtained through a single-vision lens (SVL).

All MC spectacle lenses considered here have a central clear aperture free of microstructures, with diameters ranging from 4 to 10 mm. We refer to the microstructured region outside it as the treatment area.

We assessed PSFs for several lenses and retinal defocus offsets under an indoor near-vision condition: source proximity 3 D, accommodative lag 0.5 D, and a 5 mm entrance pupil (Section 4.4.2.2). Two gaze directions were used: straight-ahead gaze and lowered gaze (20°). We varied retinal eccentricity from 0° to 30° by changing the horizontal field angle of the source. Straight-ahead gaze allowed us to assess how the projected pupil moved from the central clear aperture into the treatment area as retinal eccentricity increased. Lowered gaze (20°) was selected as a representative reading configuration in children, consistent with eye-gaze angles of approximately 20° measured during printed reading and writing tasks (Bhandari & Ostrin, 2022). At this gaze angle, the projected pupil was entirely within the treatment area at every eccentricity tested. Retinal defocus offsets were sampled from −0.5 to +0.5 D around the fixed accommodated state (Section 4.4.3).

The lenses had distinct microstructure designs and therefore produced distinct PSFs at the retina. The complete on-axis and peripheral comparisons are shown in Figure 9 and Figure 10. Notably, PLARI and NLARI used lenslets with opposite spherical powers but produced similar spatial distributions of LSC. This similarity underscores the need to go beyond geometric-optics interpretations based on lenslet power and consider the retinal image formed by the complete spectacle lens-eye system when explaining myopia-control efficacy.

### 2.2 Extracting features from indoor scenes seen through the MC spectacle lenses

We simulated how indoor scenes were imaged on the retina through each MC spectacle lens and the eye optics. To do this, we sampled RGB images, converted them to 31-wavelength hyperspectral images (HSIs), convolved each HSI with the corresponding PSFs, and projected the retinal images to grayscale using the photopic sensitivity function.

Retinal images differed across MC spectacle lens designs, each producing a distinct spatial distribution of local contrast that reflected its microstructure geometry (Figure 1, panel C).

**Figure 1.**
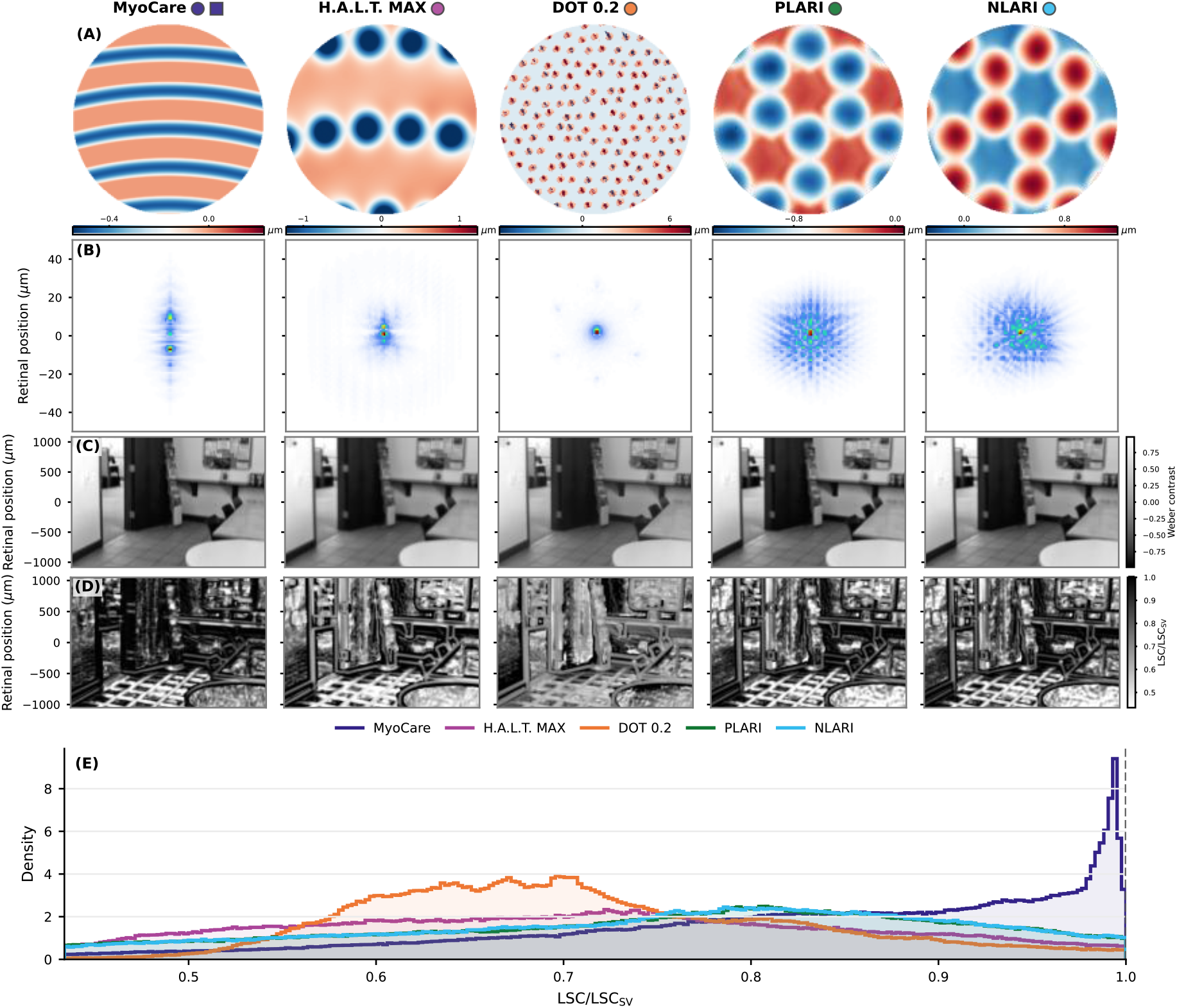
Simulation pipeline for five MC spectacle lenses, shown by column: MyoCare, H.A.L.T. MAX, DOT 0.2, PLARI, and NLARI. Symbols beside each lens name match those in Figure 2, panel A. Panels show (A) MC-layer surface sag, where positive and negative residual sag denote surface depressions and protrusions, respectively, relative to the base surface, (B) the V(λ)-weighted polychromatic PSF at the retina, (C) the Weber contrast of a V(λ)-projected retinal image from an indoor DIODE scene, (D) the corresponding LSC map divided pixelwise by the SVL map, labelled LSC/LSC_SV_, and (E) the distribution of these relative LSC values across retinal positions for each MC spectacle lens over the same displayed range as panel D. To make the retinal-image comparison easier, this illustration uses 0 D accommodative lag. The LSC maps use a receptive-field size of 15 µm.

The retina presumably extracts one or more features from the retinal image to determine the sign of defocus. In a previous study in the mouse retina, we showed that LSC carried information about defocus sign and hypothesized that the retina could use this cue to modulate eye growth (Goethals et al., 2025). Our hypothesis was that a decrease of LSC would signal that the focal plane is in front of the retina, and our prediction was that a lower LSC would be more effective at slowing down eye growth. Here, we extracted LSC from the simulated retinal images. Different MC spectacle lenses yielded distinct spatial distributions of LSC relative to the SVL (Figure 1, panels D-E), supporting LSC as a practical metric for differentiating lens designs.

### 2.3 Local spatial contrast is associated with reported clinical efficacy

LSC is the Gaussian-weighted standard deviation of Weber contrast within a retinal receptive field (RF) centered at each retinal position. RF size determines the spatial extent over which contrast is integrated. For each RF size, we calculated LSC as the root mean square of its values across retinal positions and scenes, then averaged it across the sampled retinal defocus offsets. We normalized this value to the corresponding SVL value and tested its association with reported 12-month cumulative absolute reduction in axial elongation (CARE) (Brennan et al., 2021) across RF sizes and eccentricities. Based on our previous work, we expected lower LSC relative to the SVL to be associated with greater CARE.

Under lowered gaze (20°), lower LSC was associated with greater reported CARE across a broad range of RF sizes at central and near-peripheral eccentricities, with the strongest associations at intermediate RF sizes (Figure 2, panel B). At the condition shown in Figure 2, panel A, with an eccentricity of 10° and an RF size of *σ* = 15 *µ*m, the association had *R*^2^ = 0.83 (*p* < 0.001, permutation test). Leave-one-out cross-validation of this regression gave 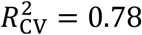 and a cross-validated root mean square error (RMSE) of 0.02 mm (Appendix, Figure 11). With straight-ahead gaze, *R*^2^ increased from central to mid-peripheral eccentricities, peaked around 20°, and then decreased at larger eccentricities (Figure 2, panel C).

**Figure 2.**
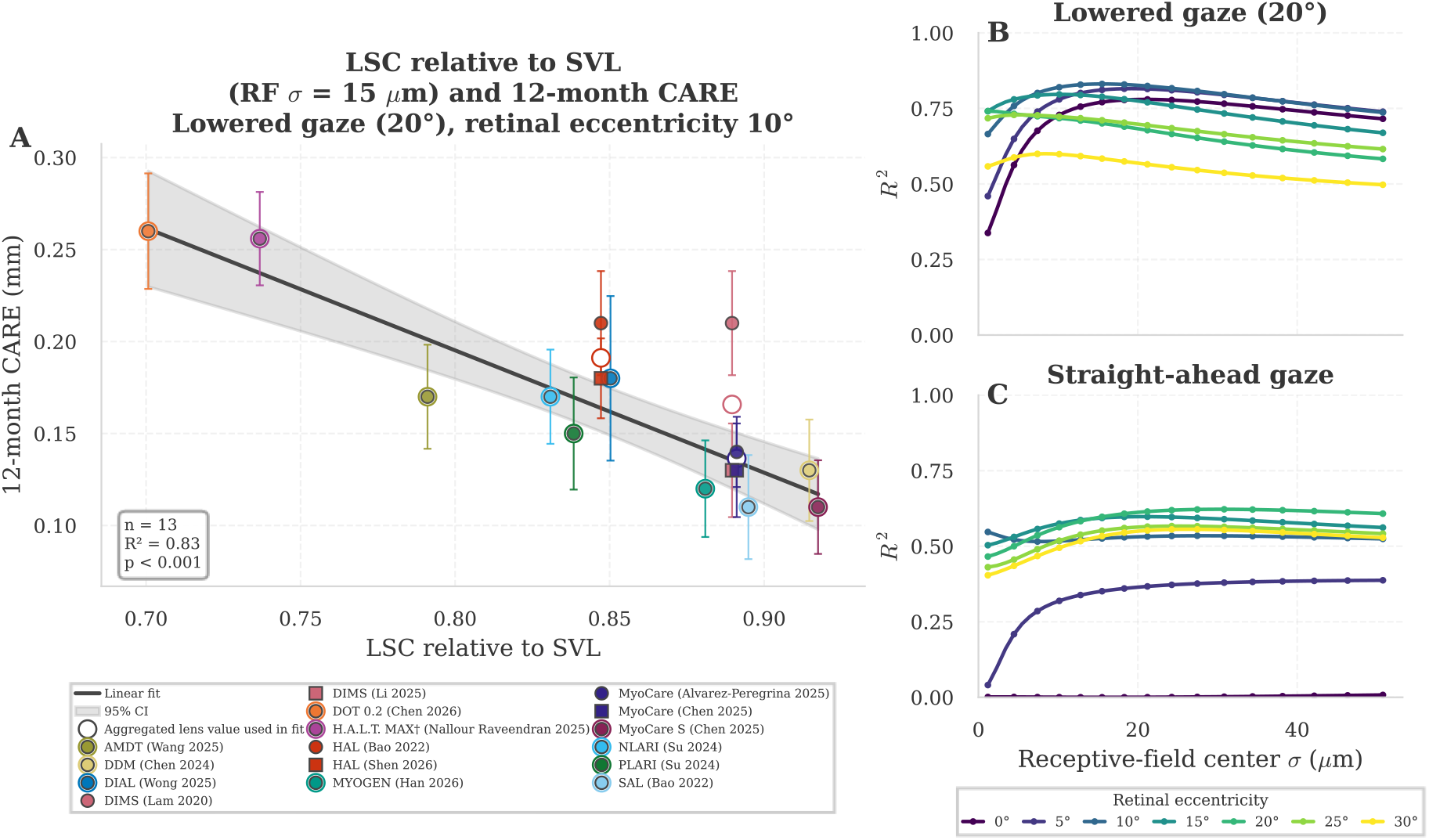
Association between LSC and reported 12-month CARE across RF sizes, retinal eccentricities, and gaze directions. (A) Association under lowered gaze (20°) at an eccentricity of 10^°^, with an RF size of σ = 15 µm. Small symbols show the individual study estimates, with vertical error bars showing their standard errors; studies of the same lens design share its simulated LSC value. Small white-filled circles show the aggregated CARE values for the 13 lens designs used in the least-squares fit, with each design contributing equally. The line shows this fit, and the shaded region shows its 95% confidence interval. (B) R^2^ as a function of RF size under lowered gaze (20°). Curves represent eccentricities from 0^°^ to 30^°^ in 5^°^ increments. (C) Corresponding R^2^ curves under straight-ahead gaze. † CARE was computed against a virtual control group (Section 4.3).

## 3. Discussion

MC spectacle lenses rely on different optical principles and have lacked a common basis for comparison. We characterized the optical impact of each design by simulating the resulting PSFs at the retina, which differed clearly across designs. We then extracted image-based metrics from the simulated retinal images and found that they were associated with reported clinical efficacy, supporting a common optical basis for comparing MC spectacle lenses. PLARI and NLARI illustrate this point: although their lenslets have opposite powers, they produced similar LSC and showed similar efficacy in the same clinical trial (CARE of 0.15 and 0.17 mm, respectively; Figure 2, panel A).

LSC showed its strongest association with reported CARE at intermediate RF sizes (Figure 2). In the frequency decomposition, increasing RF size progressively shifted the LSC weighting toward lower spatial frequencies (Figure 3). At *σ* = 15 *µ*m, near the peak CARE association, 75% of the weighting accumulated below 8.3 cpd. This overlaps the 2-8 cpd range reported to be sufficient for short-term responses to defocus, although the relative importance of frequencies within this range remains unknown (Schaeffel & Swiatczak, 2024).

**Figure 3.**
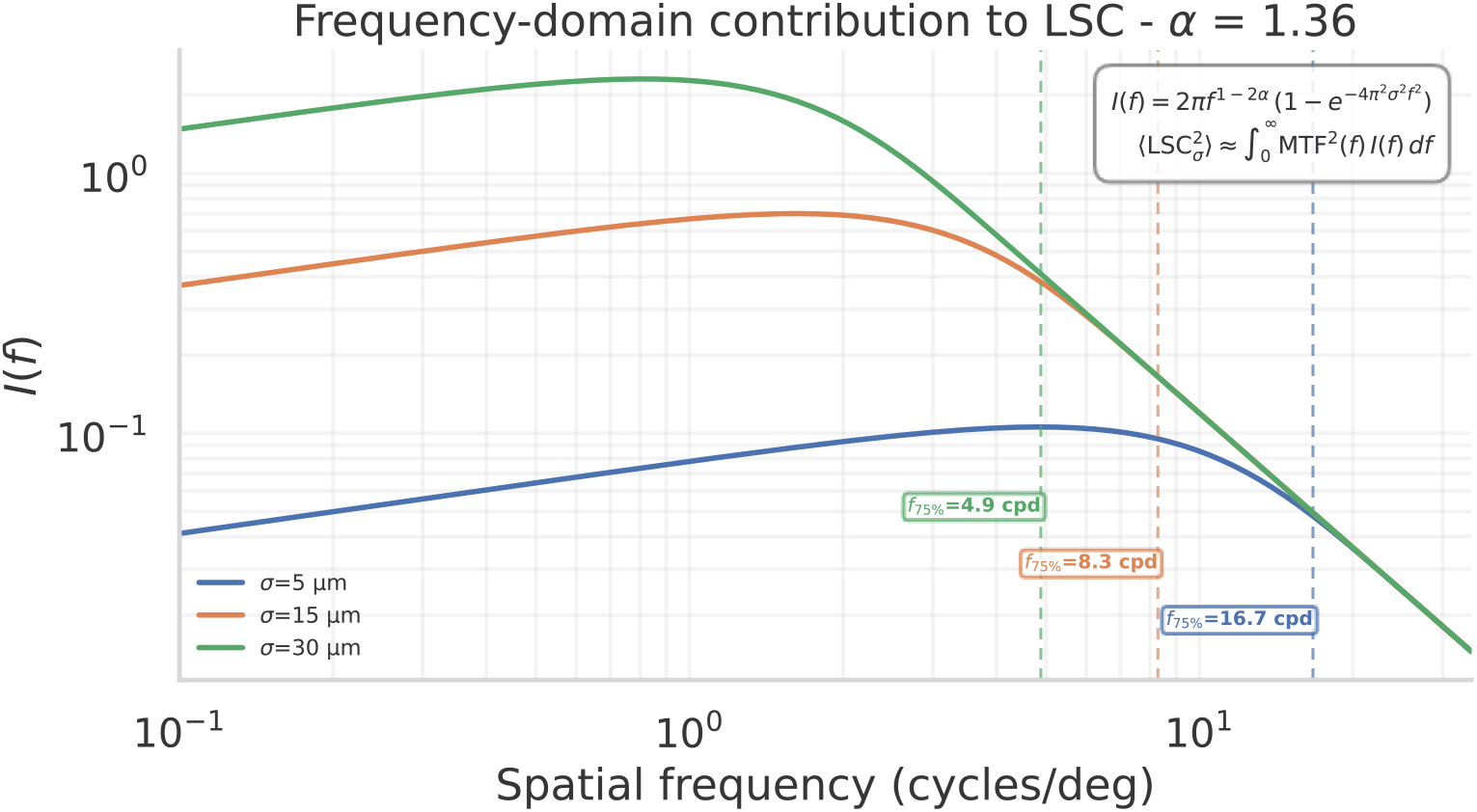
Frequency-domain decomposition of LSC for three RF sizes (5, 15, and 30 µm), plotted on logarithmic axes. Vertical markers indicate the frequency at which 75% of the total LSC^2^ has been accumulated. This threshold is lower for larger RF sizes.

The differences between gaze conditions were consistent with the regions of the lens occupied by the projected pupil. Under straight-ahead gaze, the projected pupil was predominantly located within the central clear aperture at small eccentricities. As eccentricity increased, the pupil progressively entered the treatment area. This transition is quantified in the Appendix as the fraction of entrance-pupil rays passing through the treatment area (Section 5.3).

Under lowered gaze (20°), corresponding to a near-activity posture with the projected pupil fully within the treatment area, the association was strongest at central and near-peripheral eccentricities. The peak at 10° lies within the 6° to 12° range most responsive to blur (Panorgias et al., 2021), and at the upper edge of the 6° to 10° window most sensitive to myopic defocus (Swiatczak et al., 2024). Both are short-term measurements, whereas CARE reflects axial elongation over 12 months. The present analysis cannot determine whether the peak at 10° reflects the physiological importance of this retinal region or eccentricity-dependent changes in the retinal image produced by the spectacle lens-eye system. At larger eccentricities, the association generally weakened, possibly because peripheral ocular aberrations had a greater influence on the PSFs, reducing the relative contribution of the MC spectacle lens (Section 5.1).

The association between LSC and reported clinical efficacy does not demonstrate that the retina uses LSC to regulate eye growth. LSC could provide a cue for this process, or it could vary with another relevant property of the retinal image. Distinguishing between these possibilities will require causal studies.

### 3.1 Complementary mechanisms may explain additional variation in myopia progression

The association reported here relies on a detailed model of the optics of the MC spectacle lenses and of the human eye. Our model is phenomenological and assumes that LSC carries information about defocus sign and can be computed by several retinal cell types. Although our results show that LSC carries information about reported myopia control efficacy, several complementary mechanisms may contribute to the remaining variation in myopia progression.

First, other features can be extracted to determine the sign of defocus. It has been hypothesized that chromatic aberration could provide this information through differences between the blue and green retinal images (Gawne et al., 2022; Swiatczak & Schaeffel, 2022). The MC spectacle lenses tested here did not substantially alter the relative spectral composition of the retinal image. This may explain why an achromatic feature such as LSC was strongly associated with their reported efficacy. A lens design that affects the spectral channels differently may instead require a feature that captures chromatic information (Di Tullio et al., 2026).

Factors unrelated to estimating the sign of defocus can also influence eye growth and myopia progression. These include visual-environment factors such as light exposure and circadian rhythms (Chakraborty et al., 2018), as well as genetic factors (Voogelaar et al., 2025). They may account for part of the unexplained variability. CARE partially corrects for population-level differences by comparing treatment arms with controls, but it cannot remove every confounder: age, baseline refraction, outdoor exposure, ethnicity, lifestyle, education, compliance, and study design may still modulate axial elongation. A complete model would include LSC together with clinical and environmental variables instead of treating it as a standalone optical metric.

Finally, part of the residual variability across studies may reflect the approximations of our model. The PSF computation relies on an eye model that approximates the average human eye rather than individual eyes. In addition, the spectacle lens model intentionally ignores possible freeform back-surface effects. All designs are simulated with a spherical back surface, and only the measured or reconstructed microstructured MC layer is varied. The reported optical metrics should therefore be interpreted as metrics of the microstructured layer on a common spherical carrier, not as a complete characterization of each commercial lens as worn.

### 3.2 Mechanisms of myopia progression

Because LSC is a phenomenological metric, it does not identify the retinal mechanism that detects the sign of defocus. Several ganglion cell types could extract LSC. Some amacrine cell types involved in contrast processing could also compute it and release a peptide or other modulator that initiates a signaling cascade influencing eye growth.

More direct studies identifying which cell types causally modulate eye growth are needed to clarify how MC spectacle lenses influence eye growth. A key issue will be to distinguish mechanisms that convey sign-of-defocus-dependent information from those that influence eye growth through sign-of-defocus-independent information (e.g. light level). Strategies that address both defocus-dependent and defocus-independent mechanisms through complementary approaches may improve the trade-off between myopia control efficacy on one side and visual comfort and performance on the other.

### 3.3 Outlook

The association between LSC and reported CARE is a first step toward comparing different MC spectacle lens designs on the basis of the retinal images they produce. Prospective validation is required before this metric can be used to screen new designs or predict their clinical efficacy.

Additional variation may be captured by incorporating features such as chromatic LSC. The simulation pipeline developed for this investigation can render images in L-, M-, and S-cone projection spaces, which may carry further information about defocus sign. Other factors that do not directly affect the retinal image but still influence eye growth, such as time spent outdoors, could also be integrated.

Because eye optics differ between patients, the optimal MC spectacle lens design may also vary. Once validated, the model developed here could help identify the most suitable MC spectacle lens design for each patient, supporting personalized myopia treatment informed by detailed characterization of the eye.

## 4. Methods

We describe below the simulation framework used to relate the optical effect of MC spectacle lenses to clinical efficacy. In short, we computed retinal images of indoor scenes through a spectacle lens-eye system. We then extracted image-based metrics from these simulated retinal images and tested whether they correlated with reported clinical efficacy.

### 4.1 Definitions and Acronyms

- **MC**: myopia control.
- **SVL**: single-vision lens.
- **RX**: refractive prescription.
- **Spherical back surface**: ideal smooth spectacle-lens back surface used in all simulations. Possible freeform back-surface contributions are not included.
- **Base surface**: smooth spectacle-lens surface obtained after removing the local MC microstructures from the measured or reconstructed surface. This base surface carries the underlying refractive prescription contribution.
- **MC layer**: residual microstructured sag added on top of the base surface, after subtracting the base surface from the measured or reconstructed lens geometry.
- **PSF**: point spread function.
- **LSC**: local spatial contrast.
- **RF**: receptive field.
- **CARE**: cumulative absolute reduction in axial elongation.
- **VSMTF**: visual Strehl ratio based on the modulation transfer function.
- **Source proximity**: physical source vergence, referenced to the spectacle-lens front diopter.
- **Focus proximity**: virtual source vergence, at which the accommodation maximizes visual Strehl ratio.
- **Accommodative lag**: source proximity minus focus proximity.
- **Retinal defocus offset**: defocus introduced by shifting the plane where the retinal PSF is evaluated along the chief-ray direction while keeping the optics fixed. Negative values shift this plane away from the crystalline lens and represent a more myopic state in the analysis. Positive values shift it toward the lens and represent a more hyperopic state.

### 4.2 Considered Optical Solutions

This study focuses on microstructured MC spectacle lenses. Contact lenses, progressive-addition spectacles, and bifocal spectacles are not considered. No significant differences in accommodation response amplitude were detected with DIMS, HAL, or MyoCare lenses under the tested conditions (Lin et al., 2025), whereas progressive and bifocal designs are intended to reduce accommodative demand at near. In the relaxed eye model, the visual Strehl ratio based on the MTF (VSMTF) peaked within 0.05 D of the SVL peak for every tested microstructured layer, corresponding to one sampling step. This supported using the same accommodated eye state for every design (Section 4.4.2.2).

### 4.3 Clinical Efficacy Data

We relate the retinal optical effect of each lens to its clinical efficacy. Efficacy is quantified by CARE, defined as the 12-month axial elongation of the SVL control group minus that of the treatment group. Twelve of the 13 clinical studies include such a control group. For the H.A.L.T. MAX lens evaluated in Singapore, we created a virtual control group by fitting a quadratic model to the annual axial elongation of myopic Singaporean children reported by Rozema et al. (2019). As a function of baseline age *a* in years, the fitted annual axial elongation in mm is −0.00982 *a*^2^ + 0.1095 *a* + 0.1618. Evaluating it at the H.A.L.T. MAX group’s mean baseline age of 8.6 years gave a virtual control progression of 0.38 mm over 12 months. The H.A.L.T. MAX study used a contralateral crossover design: each child wore H.A.L.T. MAX on one eye and HAL on the other, and the lenses were swapped between eyes after 6 months. Following Nallour Raveendran et al. (2025), we took the 12-month test ΔAL as the sum of the axial elongations over the two 6-month phases. This assumes no carryover between phases and no interaction between the eyes. The study found no carryover effect.

The following table lists each MC spectacle lens and the clinical study arms from which CARE was derived, grouped by lens, and reports baseline axial length (AL) and spherical equivalent refraction (SER) where available.

**Table 1.** Clinical studies used for the CARE correlation analysis, covering 13 distinct MC spectacle lens designs evaluated across 13 clinical studies. Some studies evaluated multiple designs, and some designs were evaluated in multiple studies. MC arm columns: n = cohort size; Age = baseline mean age; SER and AL are baseline values; Test ΔAL = 12-month AL change in the test arm. Ctrl ΔAL: 12-month AL change in the SVL control arm of the same study, or in the virtual control group for H.A.L.T. MAX (Section 4.3); CARE = Ctrl ΔAL - Test ΔAL.

| Design | Study | Country | n | Age (y) | SER (D) | AL (mm) | Test ΔAL (mm) | Ctrl ΔAL (mm) | CARE (mm) |
| --- | --- | --- | --- | --- | --- | --- | --- | --- | --- |
| **AMDT** | (Wang 2025) | China | 71 | 10.4 | -2.43 | 24.56 | 0.16 | 0.33 | 0.17 |
| **DDM** | (Chen 2024) | China | 81 | 10.2 | -2.74 | 24.62 | 0.21 | 0.34 | 0.13 |
| **DIAL** | (Wong 2025) | Singapore | 38 | 10.8 | -2.50 | 24.50 | 0.04 | 0.22 | 0.18 |
| **DIMS** | (Lam 2020) | China | 79 | 10.2 | -2.97 | 24.70 | 0.11 | 0.32 | 0.21 |
|  | (Li 2025) | China | 72 | 10.7 | -3.16 | 24.82 | 0.17 | 0.30 | 0.13 |
| **DOT 0.2** | (Chen 2026) | China | 128 | 9.6 | -2.36 | 24.49 | 0.09 | 0.35 | 0.26 |
| **H.A.L.T. MAX** | (Nallour Raveendran 2025) | Singapore | 50 | 8.6 | -1.90 | 23.98 | 0.12 | 0.38 | 0.26 |
| **HAL** | (Bao 2022) | China | 54 | 10.7 | -2.70 | 24.76 | 0.14 | 0.35 | 0.21 |
|  | (Shen 2026) | USA | 77 | 9.8 | -2.21 | 24.26 | 0.09 | 0.27 | 0.18 |
| **MyoCare** | (Alvarez-Peregrina 2025) | Europe | 115 | 10.0 | -2.28 | 24.34 | 0.08 | 0.22 | 0.14 |
|  | (Chen 2025) | China | 78 | 9.9 | -2.23 | 24.44 | 0.19 | 0.32 | 0.13 |
| **MyoCare S** | (Chen 2025) | China | 78 | 9.8 | -2.30 | 24.34 | 0.21 | 0.32 | 0.11 |
| **MYOGEN** | (Han 2026) | China | 79 | 10.4 | -2.15 | 24.44 | 0.25 | 0.37 | 0.12 |
| **NLARI** | (Su 2024) | China | 77 | 9.7 | -2.26 | 24.43 | 0.17 | 0.34 | 0.17 |
| **PLARI** | (Su 2024) | China | 76 | 9.4 | -2.30 | 24.47 | 0.19 | 0.34 | 0.15 |
| **SAL** | (Bao 2022) | China | 55 | 10.1 | -2.31 | 24.43 | 0.24 | 0.35 | 0.11 |

### 4.4 Optical simulations

Below, we summarize the retinal-image simulation pipeline used for each lens. Each step is detailed in its own subsection:

1. measure or reconstruct the MC layer (Section 4.4.1);
2. derive focus proximity from source proximity and accommodative lag using the eye model equipped with an SVL, and reuse this focus proximity for all assessed MC spectacle lenses, as supported by the best-focus comparison in Section 4.4.2.2;
3. construct the spectacle lens-eye system with this accommodated state (Section 4.4.2.3);
4. compute a polychromatic, through-focus set of PSFs at the retina (Section 4.4.3);
5. sample a set of indoor RGB scenes and convert them to 31-wavelength HSIs (Section 4.4.4);
6. convolve each HSI with the set of PSFs at the corresponding wavelengths, then project the resulting images with a selected spectral sensitivity *Q*(*λ*), yielding a simulated achromatic retinal image at each retinal defocus offset (Section 4.4.5);
7. derive the LSC metric from each simulated retinal image (Section 4.5.1).

Most of the pipeline is implemented with custom software.

The simulations use an indoor near-vision condition with a 5 mm entrance pupil. Source proximity is 3 D at the spectacle-lens front diopter and accommodative lag is 0.5 D, giving a focus proximity of 2.5 D (Section 4.4.2.2). Two gaze directions are considered: lowered gaze (20°), used for the full image-based computation, and straight-ahead gaze. The source direction is swept along the horizontal meridian, with eccentricity ranging from 0° to 30°. Retinal defocus offsets are sampled from −0.5 to +0.5 D in 0.125 D increments around the fixed accommodated state (Section 4.4.3). The eye optics remain fixed. Negative values shift the retinal PSF analysis plane away from the crystalline lens along the chief-ray direction and represent a more myopic refractive state in the analysis. Positive values shift it toward the lens and represent a more hyperopic refractive state.

#### 4.4.1 Spectacle lens measurements

We measured the surface geometry of all MC spectacle lenses except AMDT, DDM, MyoCare, and MyoCare S, which we reconstructed from publicly available descriptions and manufacturer specifications, including white papers.

Lens surface measurements were performed using a NewView™ 9000 optical profiler (Zygo Corporation, Middlefield, CT, USA). Based on Coherence Scanning Interferometry (de Groot, 2015), this instrument provides non-contact three-dimensional surface measurements. Its axial resolution and accuracy are well suited to measuring microstructures manufactured on lens surfaces. The instrument is used with a 5-axis motorized stage and a controller (joystick) for X-Y, encoded Z and pitch/roll (tilt) movement. This stage allows the sample to be aligned under the microscope objective. For the characterization of the different samples, the optical system was mainly configured with a 5.5× objective, a 0.5× field zoom lens, and a camera operating at 195 Hz, providing an effective acquisition area of 1000 pixels × 1000 pixels.

As a microscope interferometer, the optical profiler provides a field of view (FOV) that depends on the optical configuration. For example, with the main optical configuration used in this study, the maximum FOV achievable is about 3.1 mm × 3.1 mm (with the current lateral calibration). To achieve a larger FOV, a stitching process can be used. First, overlapping images are acquired. This is done by translating the stage along the X and Y axes to move the lens in a controlled way under the objective. The images are then stitched with the dedicated algorithms in Mx™ software for the NewView™ 9000, yielding an extended FOV.

This FOV extension depends on the surface curvature. For a flat surface, it is limited by the translation range and the computing power to process the stitching. However, microstructured surfaces of myopia control spectacle lenses are curved. Before measuring a curved surface, pitch and roll of the motorized stage must be adjusted to align the surface normal with the objective optical axis. This alignment step is needed to optimize surface measurement. Indeed, as explained in de Groot (2015), the numerical aperture (NA) of the objective limits the collection angle of specular reflections. This limit is given in terms of range of surface slopes. A high magnification objective has a higher NA, allowing for the measurement of steeper slopes at the expense of a reduced FOV. Moreover, local slopes of the surface induce aberrations and retrace errors that impact the measurement, as studied in Shahinian et al. (2021).

Because stitching is performed with simple X-Y translation, pitch and roll are not re-adjusted at each measurement site to keep the local surface normal to the objective optical axis. Therefore, the curvature limits the FOV extension. In our current configuration, the largest achievable extended FOV is about 10 mm × 10 mm for a surface with a radius of curvature of about 168 mm. Other factors also limit this extension, such as anti-reflective coatings that significantly reduce the amount of reflected light.

To overcome the limited extension of FOV on myopia control spectacle lenses with curved surfaces, we developed an internal stitching method based on circular movement of the lens. Instead of moving the test lens only along the X and Y axes with the motorized stage, we combined this stage with an additional 360° rotary stage (*θ* axis). The new method uses circular stitching patterns. After aligning the center of rotation of both the stage and the lens under test, a starting point was defined at a fixed radial distance for the first measurement site. For this starting point, pitch and roll were adapted to make the surface normal to the objective optical axis. Then, an annular stitching pattern was performed by changing the *θ* axis to rotate the lens over 360°. The angular step depends on the distance from the center and must be small enough to overlap images. Because the X-Y position of each site does not physically change while *θ* does, each site’s spatial coordinates were recalculated from the stage rotation angles and the starting-point (*x, y*) coordinates, with (0,0) defined at the lens center. Finally, the same stitching algorithms in Mx™ software were used to stitch all measurements together. Different starting points can be defined at different radial positions so the complete lens surface can be measured. Numerous precautions are needed to minimize measurement errors associated with this method. Their description is beyond the scope of the present discussion. Each measured or reconstructed lens surface is decomposed into a base surface and an *MC layer* ℒ = (*z*_MC_(*x, y*), Δ*n*). The base surface is the smooth residual surface after the microstructures have been removed. Only the low-order shape remains, which is used for the ametropia correction. The function *z*_MC_(*x, y*) is the residual sag of the MC layer relative to that base surface. The term Δ*n* is the refractive-index difference between the lens material and air at the design wavelength.

The surface sag profiles of all considered microstructured MC layers are shown in Figure 4.

**Figure 4.**
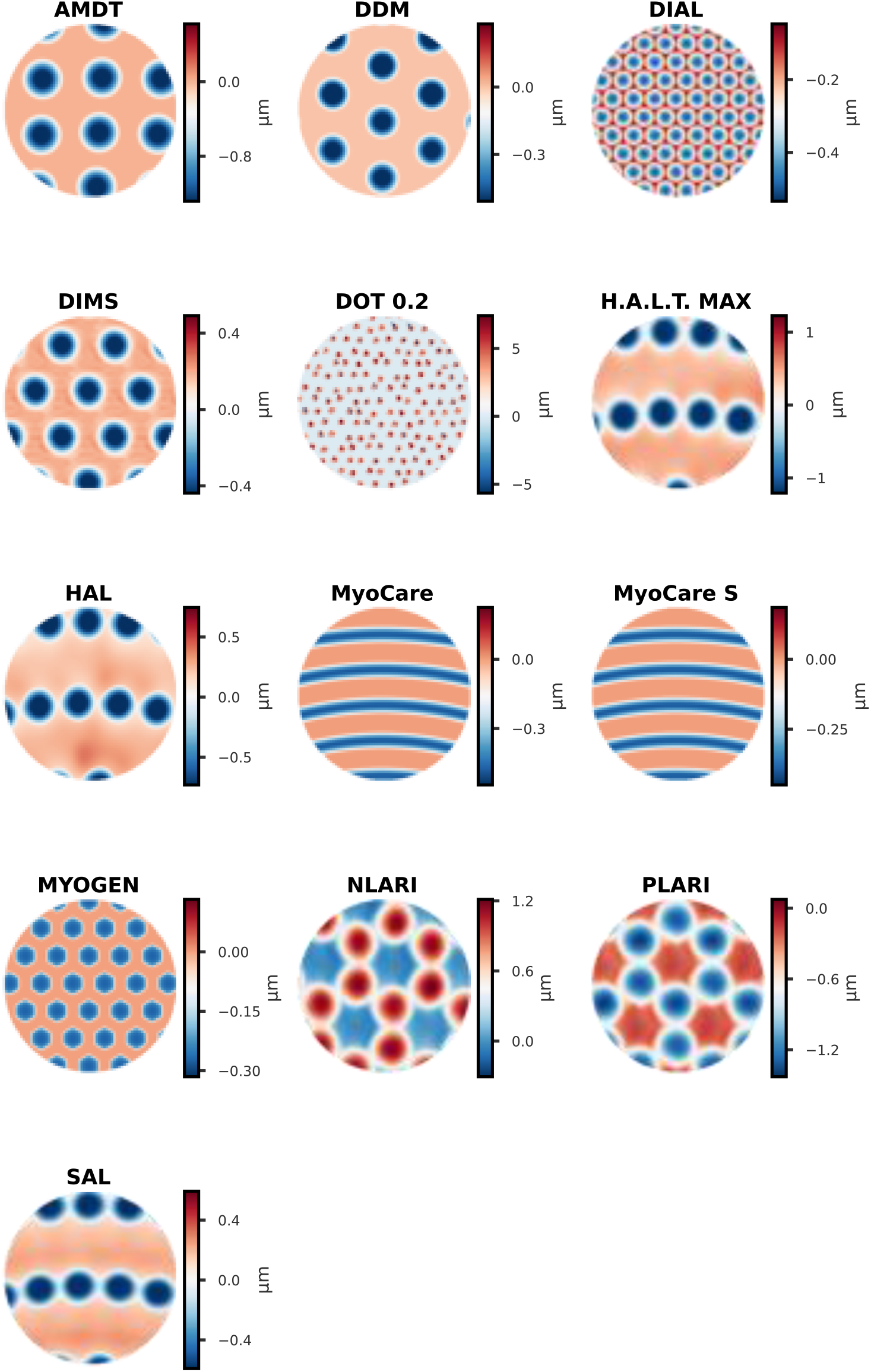
The surfaces correspond to a 5 mm entrance pupil with lowered gaze (20°) and an on-axis source. The sign convention is the same as in Figure 7.

Importantly, this decomposition is used to isolate the contribution of the microstructured layer. We do not model possible freeform back surfaces of the commercial lenses. In all simulations, the spectacle lens back surface is replaced by a spherical surface compatible with the −2 D refractive prescription (RX).

#### 4.4.2 Spectacle lens-eye system

##### 4.4.2.1 Relaxed eye model

We use a representative −2 D modified myopic eye model based on centered Model 1 from Atchison (2006). The model was adjusted from the original geometry with the following objectives while maintaining a spherical equivalent refractive error of −2 D:

1. **Peripheral refraction**. Peripheral mean sphere (*M*) data from Sng et al. (2011), Li et al. (2015) (horizontal meridian) and Chen et al. (2010) (vertical meridian) were aggregated, then interpolated for a −2 D central equivalent mean sphere.
2. **Primary spherical aberration**. The relaxed eye model was adjusted using measurements reported by Hiraoka et al. (2017) and Lau et al. (2018), accounting for differences in pupil diameter by changing the model pupil. At a 5 mm entrance pupil, the resulting target was 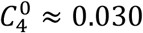 *µ*m.
3. **Peripheral astigmatism**. The original Atchison model overestimates peripheral astigmatism relative to population norms. We used peripheral astigmatism measurements in emmetropic eyes from Gustafsson et al. (2001) as a target.

The resulting surface parameters are listed in Table 2. All ocular media follow a Cauchy dispersion model *n*(*λ*) = *A* + *B*/*λ*^2^ (with *λ* in *µ*m) so that chromatic effects are captured directly during ray tracing.

**Table 2.**
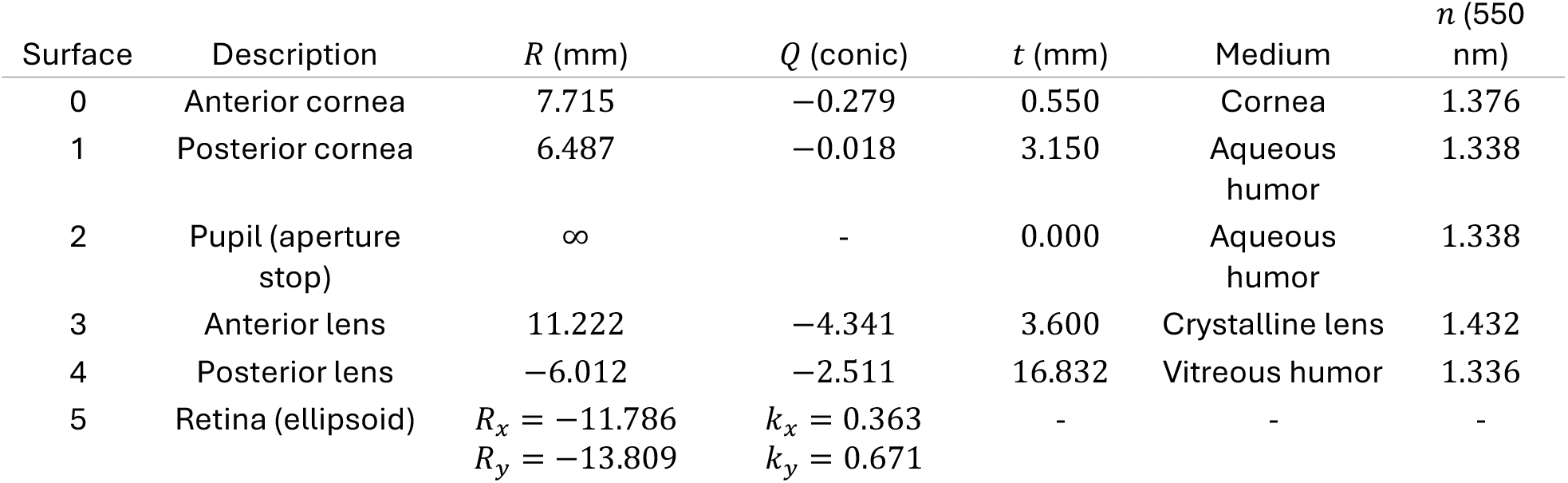
Relaxed eye-model surface parameters. R = radius of curvature; Q = conic constant; t = axial distance to the next surface. Retina is a biconic ellipsoid with principal radii R_x_, R_y_ and conic constants k_x_, k_y_. Refractive indices n are evaluated at λ = 550 nm from the Cauchy law.

| Surface | Description | $R$ (mm) | $Q$ (conic) | $t$ (mm) | Medium | $n$ (550 nm) |
| --- | --- | --- | --- | --- | --- | --- |
| 0 | Anterior cornea | 7.715 | $-0.279$ | 0.550 | Cornea | 1.376 |
| 1 | Posterior cornea | 6.487 | $-0.018$ | 3.150 | Aqueous humor | 1.338 |
| 2 | Pupil (aperture stop) | ∞ | - | 0.000 | Aqueous humor | 1.338 |
| 3 | Anterior lens | 11.222 | $-4.341$ | 3.600 | Crystalline lens | 1.432 |
| 4 | Posterior lens | $-6.012$ | $-2.511$ | 16.832 | Vitreous humor | 1.336 |
| 5 | Retina (ellipsoid) | $R_x = -11.786$<br>$R_y = -13.809$ | $k_x = 0.363$<br>$k_y = 0.671$ | - | - | - |

Dispersion coefficients: cornea (*A* = 1.3760, *B* = 8.84 × 10^−6^ *µ*m^2^), aqueous humor (*A* = 1.3338, *B* = 1.12 × 10^−3^ *µ*m^2^), crystalline lens (*A* = 1.4242, *B* = 2.33 × 10^−3^ *µ*m^2^), vitreous humor (*A* = 1.3359, *B* = 3.50 × 10^−5^ *µ*m^2^). The resulting LCA spans approximately −1.95 D at 380 nm to +0.80 D at 780 nm, consistent with published ocular LCA data.

These parameters define the relaxed eye model from which accommodated states are derived.

##### 4.4.2.2 Accommodation model

Source proximity *p*_*s*_ describes the physical source vergence relative to the spectacle-lens front diopter. Focus proximity *p*_*f*_ describes the modeled accommodative response, and accommodative lag is defined as *p*_*s*_ − *p*_*f*_. For the principal near condition, *p*_*s*_ = 3 D and the assumed lag is 0.5 D, giving *p*_*f*_ = 2.5 D. Kaphle et al. (2022) reported an accommodative lag of 0.59 ± 0.07 D in myopic eyes using a visual Strehl ratio metric and natural pupils, supporting the selected approximate value.

Starting from the relaxed eye model, we optimized crystalline-lens shape to maximize on-axis foveal VSMTF through the SVL while matching the modeled accommodation-induced changes in primary and secondary spherical aberration, 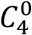 and 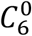, respectively. The VSMTF uses the contrast sensitivity function of Rovamo et al. (1978) at 550 nm with a 5 mm entrance pupil. To construct an eye state at focus proximity *p*_*f*_, the optimization evaluated best focus using a reference source at *p*_*f*_. The resulting eye geometry is then fixed, and retinal PSFs are computed at the physical source proximity *p*_*s*_, yielding slightly defocused images.

The optimization has three independent degrees of freedom:

1. additional anterior crystalline-lens surface power Δ*P*_anterior_, with the posterior power increment set to half the anterior increment, reflecting its smaller change during accommodation (Navarro, 2014);
2. the anterior crystalline-lens conic constant;
3. the posterior crystalline-lens conic constant.

The targets for the accommodation-induced changes in 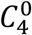 and 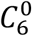 are taken from the closed-form model of Lopez-Gil & Fernandez-Sanchez (2010). With increasing focus proximity, the resulting 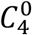 decreases, whereas 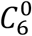 remains positive and increases (Figure 5).

**Figure 5.**
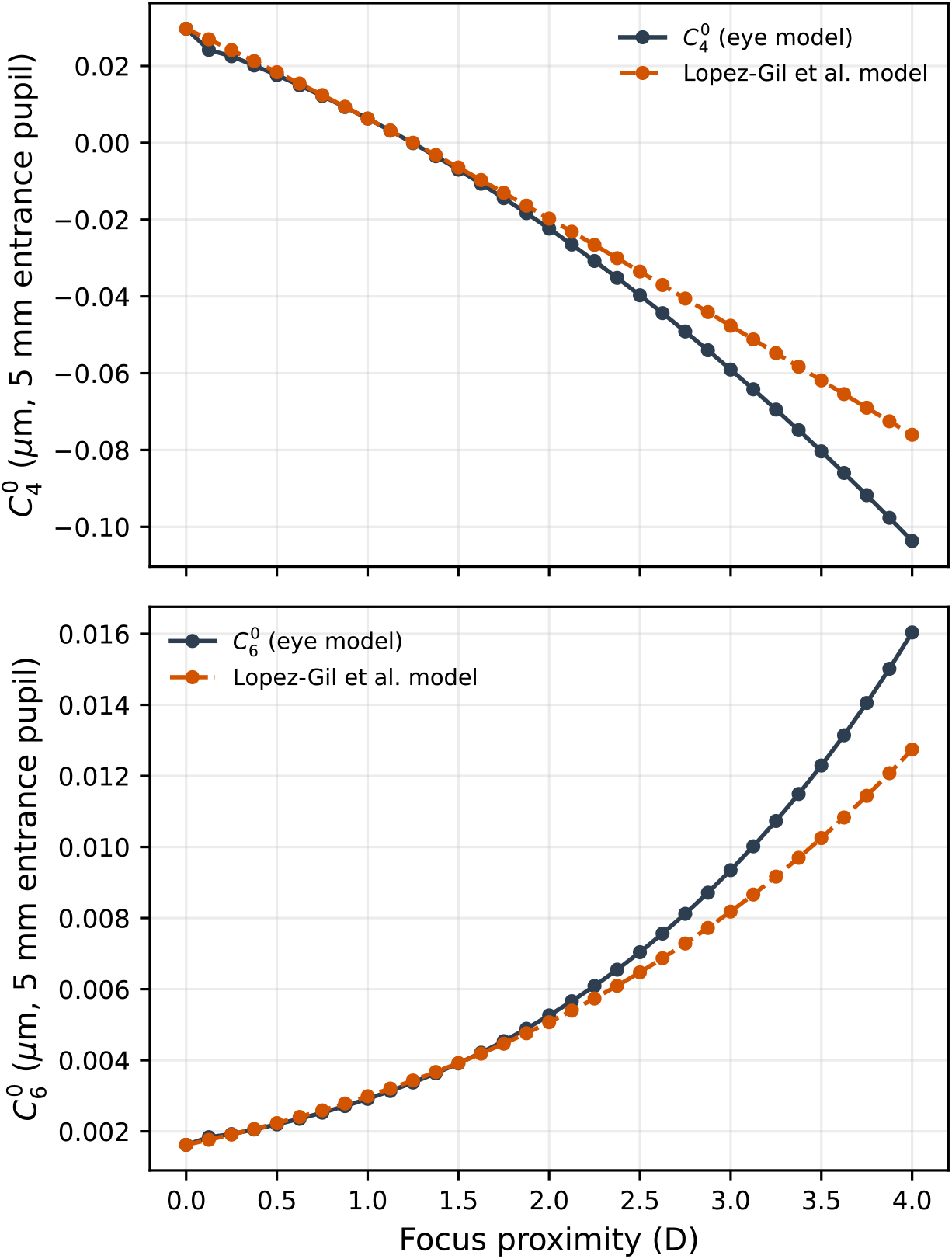
Primary and secondary spherical-aberration coefficients for the representative −2 D eye model as a function of focus proximity from 0 to 4 D. The upper panel shows 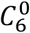, and the lower panel shows 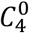. Dark solid lines show the eye-model coefficients. Orange dashed lines show the accommodation-induced changes from Lopez-Gil & Fernandez-Sanchez (2010), added to the corresponding relaxed-eye coefficient at 0 D. Values are calculated for straight-ahead gaze at 550 nm with a 5 mm entrance pupil. Accommodated states maximize foveal VSMTF through the SVL while targeting the modeled changes in both spherical-aberration coefficients.

Accommodation microfluctuations continually perturb retinal defocus (Charman & Heron, 2015). We therefore sample retinal defocus offsets over ±0.5 D around one fixed accommodated eye state rather than evaluating only one value.

We also tested whether the microstructured layers shifted modeled optical best focus relative to the SVL. For this validation, the relaxed eye viewed an on-axis source at 2.5 D, matching the focus proximity of the principal near condition, through each lens. Source proximity was referenced to the spectacle-lens front diopter, and the simulation used lowered gaze (20°). VSMTF was evaluated at 550 nm with a 5 mm entrance pupil while the retinal PSF analysis plane was shifted along the chief-ray direction over defocus offsets from −3.0 to −1.50 D in 0.05 D increments. Best-focus offsets differed from that of the SVL by at most 0.05 D (Figure 6).

**Figure 6.**
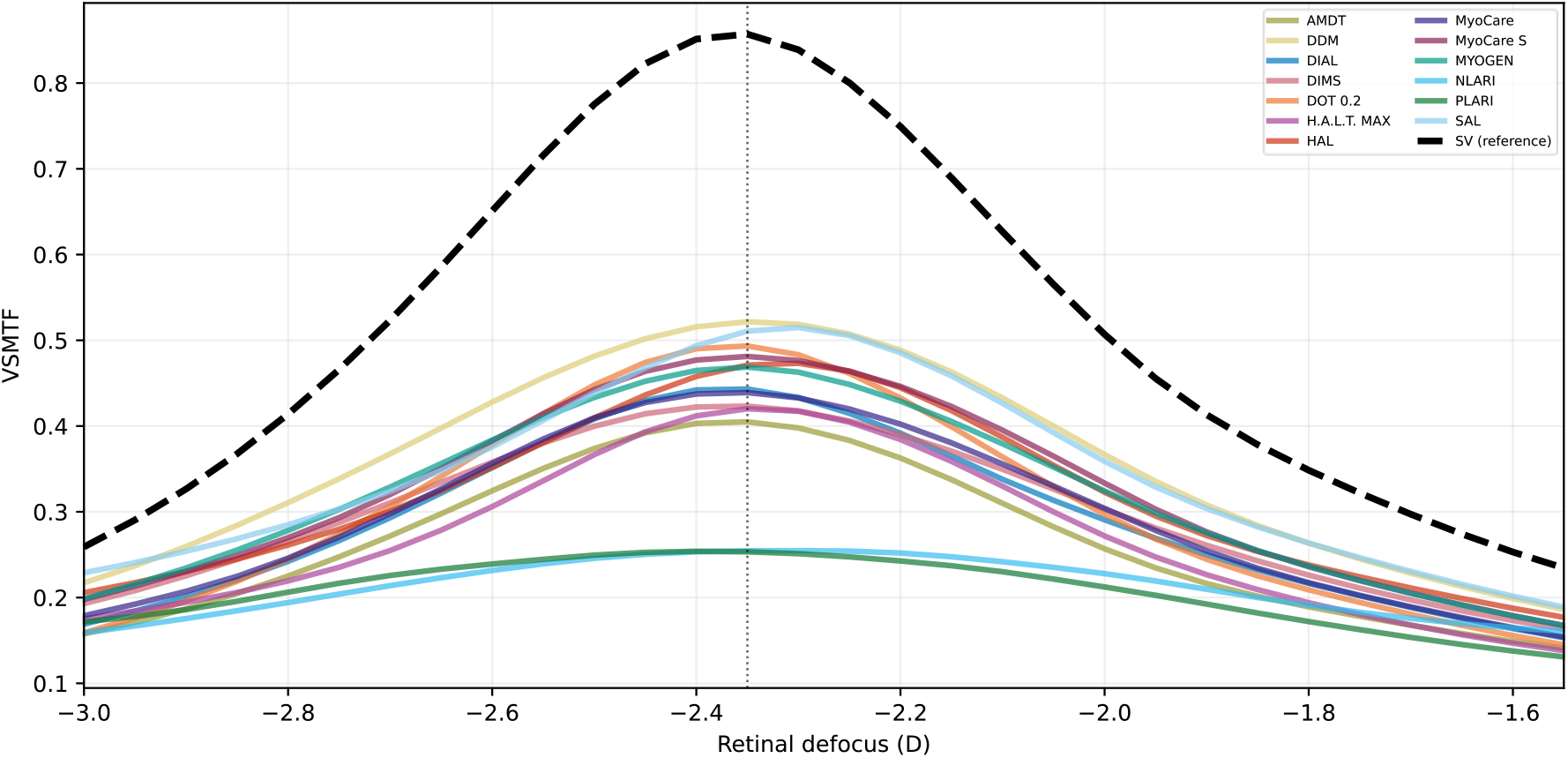
Through-focus VSMTF for the tested MC spectacle lenses (colored lines) and the SVL (dashed black). The relaxed-eye simulation uses a 2.5 D source proximity at the spectacle-lens front diopter, lowered gaze (20°), a 5 mm entrance pupil, and a wavelength of 550 nm. Peak offsets within 0.05 D of the SVL peak support calculating accommodation with the SVL and using the resulting eye state for every tested microstructured layer.

We therefore determine accommodation once with the SVL and reuse that accommodated eye state for every tested MC spectacle lens at a given source proximity.

##### 4.4.2.3 Spectacle lens-eye construction

We construct the spectacle lens-eye system for a given MC layer in four steps:

- a smooth base surface corresponding to a standard SVL is created with power −2 D and refractive index *n*_ℒ_ = 1 + Δ*n*, matching the material of the MC layer;
- the back surface of this SVL carrier is kept spherical for all designs;
- the MC layer *z*_MC_(*x, y*) is added to that base surface, so that the full front surface becomes *z*_front_(*x, y*) = *z*_base_(*x, y*) + *z*_MC_(*x, y*);
- the eye model is fitted with the reconstructed spectacle lens, and the eye is oriented along the prescribed gaze direction (*α, β*).

The figure below illustrates the propagation geometry used in the simulations.

**Figure 7.**
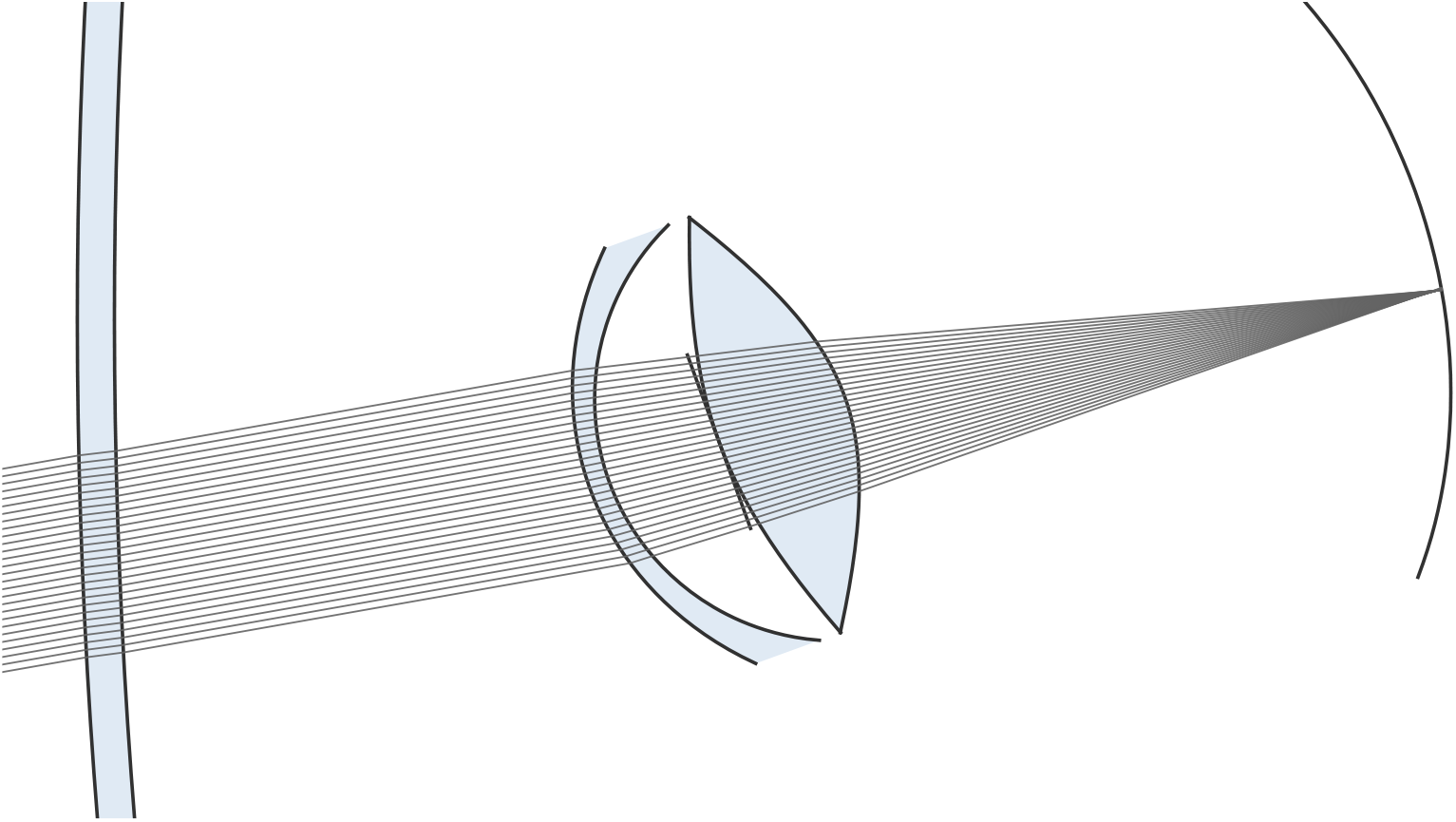
Propagation geometry used to simulate retinal image formation through an MC spectacle lens.

#### 4.4.3 Polychromatic PSFs

For an eye focused at a proximity of 2.5 D, we kept the spectacle lens-eye optics fixed and calculated monochromatic PSFs at retinal defocus offsets from −0.5 to +0.5 D in 0.125 D increments.

We denote by 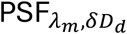 (*i, j*) the monochromatic PSF at wavelength *λ*_*m*_ and retinal defocus offset *δD*_*d*_.

For each wavelength and retinal defocus offset (*λ*_*m*_, *δD*_*d*_), with *δD*_*d*_ expressed in diopters, we proceed as follows:

- the retinal defocus offset *δD*_*d*_ is converted to a signed displacement Δ*s*_*d*_ =
- −0.341 mm/D × *δD*_*d*_ of the PSF analysis plane along the chief-ray direction. Positive Δ*s*_*d*_ is directed away from the crystalline lens. The conversion coefficient was calibrated by relating trial-lens power to the displacement along the chief ray required to recover best focus.
- a source beam is defined in object space so that it reaches the cornea with the desired eccentricity (*h, v*) after traversing the spectacle lens.
- the source beam is sampled on a uniform 1024 × 1024 ray grid across the entrance pupil and propagated to the system exit pupil while accumulating optical path length.
- the Stiles-Crawford effect was included by applying the apodization function exp(−*ρ*_*e*_*r*^2^/2), with *ρ*_*e*_ = 0.10592 mm^−2^, thus reducing the importance of peripheral pupil rays (Zhang et al., 1999).
- the complex pupil function is built from the optical path difference sampling, from which we obtain the Fraunhofer PSF on the retina using non-uniform fast Fourier transform (NUFFT) integration. We use the Type 3 NUFFT implemented in the FINUFFT/cuFINUFFT library (Shih et al., 2021).

This final step matters because MC spectacle lenses transform an initially uniform ray grid into a non-uniform distribution. The Type 3 NUFFT lets us integrate from these non-uniform exit-pupil points directly onto the uniformly sampled retinal image grid. The PSF is evaluated on a 1024 × 1024 retinal grid covering approximately 1 mm^2^, i.e. an initial sampling of about 1 *µ*m/pixel.

#### 4.4.4 RGB images to HSIs

Real-world scenes vary in terms of their spectral composition and the ocular media are dispersive, so retinal image formation is wavelength-dependent. Ideally, we would use a large HIS database and combine it with the respective L-, M-, and S-cone spectral sensitivities and spatial organization. However, such datasets remain scarce. Most large scene datasets contain only RGB images. Furthermore, interference and diffractive effects become particularly pronounced in monochromatic PSFs, making a simple three-wavelength RGB simulation inadequate. HSIs were therefore generated from the RGB DIODE dataset (Vasiljevic et al., 2019), which contains more than 25 000 indoor and urban-outdoor images across 27 scenes. We retained only the indoor subset for our simulations. Indoor near-distance scenes were selected to represent conditions relevant for MC simulations: near-work behavior, including shorter viewing distance and prolonged near work, has been associated with faster myopia progression (Huang et al., 2020), and indoor environments have spatial-frequency characteristics distinct from natural outdoor scenes and potentially relevant to myopia development (Flitcroft et al., 2020).

DIODE also provides a high-precision depth map (±1 mm) for each image, but we do not use it here. Object distance has a direct effect on retinal image formation: objects at the accommodated distance form the sharpest retinal image, closer objects focus behind the retina, and farther objects focus in front of it. We ignored this effect in the present simulations. Including depth would require depth-dependent PSF simulations and is left for future work.

We denote by *S*_RGB_(*i, j, c*) a source RGB image from DIODE, where *c* indexes the RGB color channel. We estimate the sampled HIS *S*(*i, j, λ*_*m*_) from *S*_RGB_ using MST++ (Cai et al., 2022), a transformer trained for RGB-to-HIS reconstruction on the ARAD 1K dataset used in the NTIRE 2022 Spectral Reconstruction Challenge. MST++ outputs 31 wavelengths from 400 to 700 nm in 10 nm steps. Note that MST++ is trained on a limited set of scenes (900 HIS-RGB pairs). It may therefore introduce wavelength-dependent biases outside its training distribution. We interpret the reconstructed spectrum as a plausible spectral distribution consistent with the RGB image, not as a radiometric measurement.

We extracted a representative subset of *N*_img_ = 150 indoor scenes from DIODE. We compare the spatial-frequency statistics of this subset with values reported for indoor images by Flitcroft et al. (2020) and D.-L. Li et al. (2025).

Images in the DIODE dataset have resolution 1024×768 pixels. Their nominal field of view is 60°×45°, which would place the Nyquist frequency at only 8.5 cpd. Since our simulations cover both foveal and peripheral vision, and the foveal contrast sensitivity function extends well over 30 cpd (Rovamo et al., 1978), this sampling is insufficient to represent the foveal spatial-frequency range. We therefore treat the images as covering a reduced field of 15°×11.25°, raising the Nyquist frequency to 34.1 cpd.

The spatial-frequency content of the selected indoor subset is shown below. The log-log slope was fitted over the (0.1) to (10) cpd range, consistent with the fitting range used in the reference studies cited above.

The log-log slope of the indoor subset was comparable to published values for indoor scenes.

**Figure 8.**
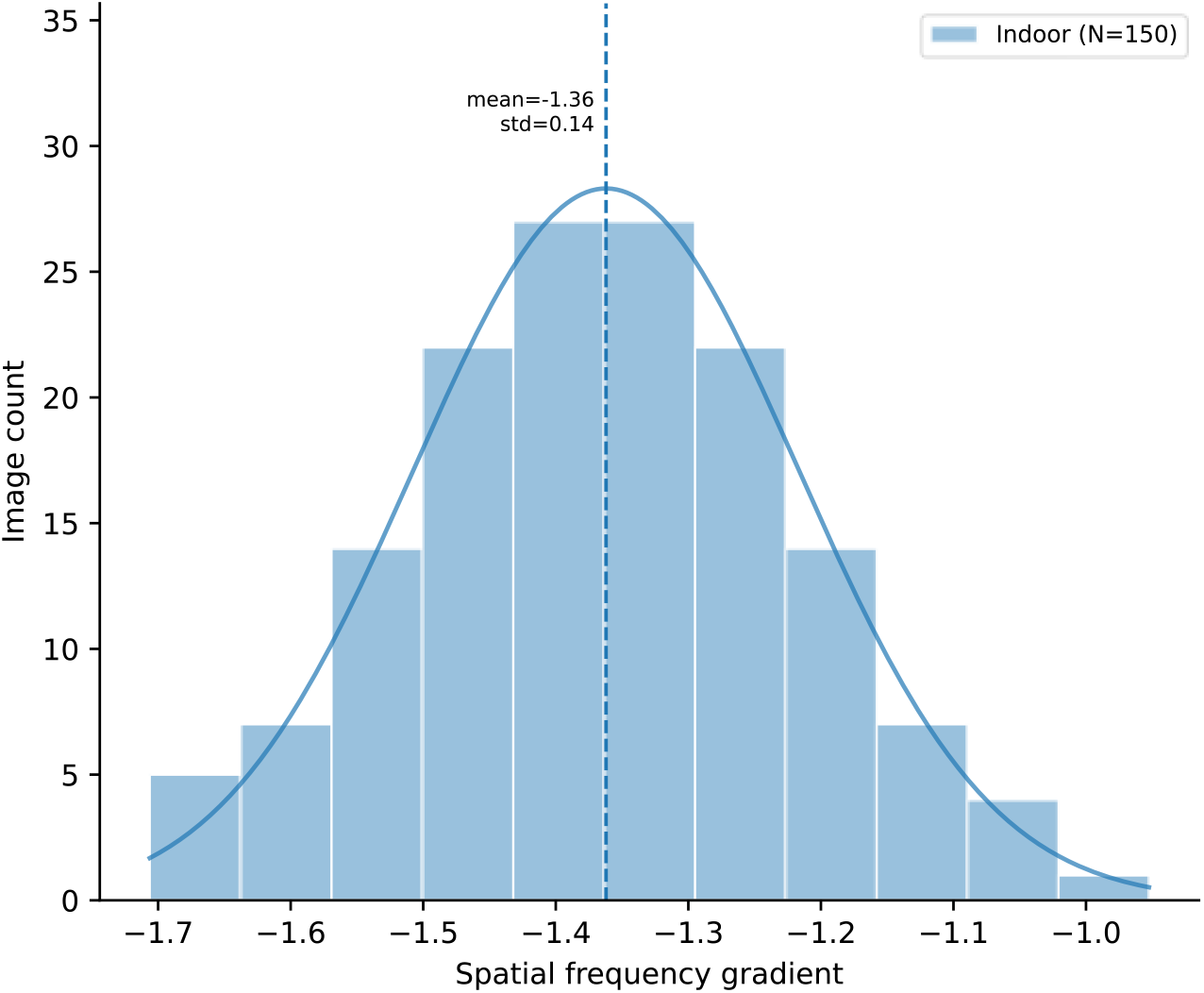

##### 4.4.4.1 Image preprocessing

Before convolution, each reconstructed HSI is normalized in the same projection space *Q*(*λ*) used for retinal image simulation (Section 4.4.5). Let

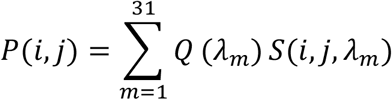

be the projected image. If *µ*_*P*_ and *σ*_*P*_ are the spatial mean and standard deviation of *P*, we define

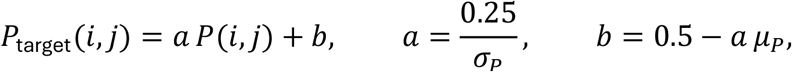

so that the target projected image has a mean projected signal of 0.5 and a standard deviation of 0.25. The HSI is then normalized as

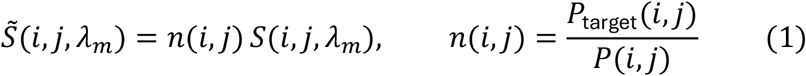

The same scaling factor is therefore applied to all wavelengths at a given pixel. This standardizes global luminance and contrast across scenes, so that LSC is less affected by global brightness or contrast differences between images. Similar normalization of real-world image sets before LSC analysis is used by Karamanlis & Gollisch (2021) and Liu et al. (2022) for grayscale images. Here, we extend this approach to HSIs.

#### 4.4.5 Retinal image simulation

We now reconstruct the retinal image. Let *S*(*i, j, λ*_*m*_) be the source HSI (Section 4.4.4) and 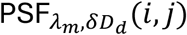 (*i, j*) the PSF from Section 4.4.3, normalized so that 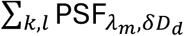 (*k, l*) = 1.

The source images and initially generated 1024 × 1024 PSFs are resampled to a common angular sampling of 1/120 degree per pixel, giving a Nyquist frequency of 60 cycles per degree. This yields a 1350 × 1800 image grid and 420 × 420 PSFs for convolution. The monochromatic retinal image at wavelength *λ*_*m*_ and retinal defocus offset *δD*_*d*_ is

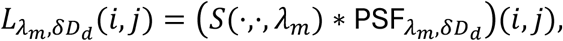

where *\** denotes 2-D spatial convolution, computed in the frequency domain via a fast Fourier transform (FFT) with valid-region padding to avoid border artifacts.

We then project across wavelengths with a selected spectral sensitivity *Q*(*λ*):

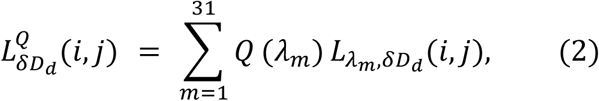

Here *Q*(*λ*) is chosen as the International Commission on Illumination (CIE) 1924 photopic luminosity function *V*(*λ*), a pooled long- and medium-wavelength cone sensitivity *LM*(*λ*), or the short-wavelength cone sensitivity *S*(*λ*). This yields one projected retinal image for each retinal defocus offset and each selected spectral sensitivity. The photopic projection captures luminance, whereas cone-based projections support chromatic comparisons.

In short, each source image is processed wavelength by wavelength. We convolve the spectral scene with the corresponding spectral PSF, then project the result with a chosen spectral sensitivity.

### 4.5 Metric Calculations

#### 4.5.1 Local spatial contrast

We selected LSC as a key retinal-image metric because it provides a spatially local measure of contrast after optical transformation by the spectacle lens-eye system. This choice was motivated by previous work in a mouse eye model suggesting that LSC may contribute to retinal estimation of defocus sign and visually guided eye-growth regulation (Goethals et al., 2025). Although that work was performed in a mouse model and cannot be assumed to transfer directly to humans, it provides a mechanistic basis for testing whether such an image-based contrast metric is associated with reported clinical efficacy in humans.

LSC quantifies how much local contrast a small retinal region contains: for each retinal position, we weight neighboring pixels by an RF profile and measure how much contrast varies within that RF.

We follow the definition described in Karamanlis & Gollisch (2021). For a given retinal defocus offset *δD*_*d*_, the projected retinal image 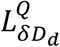 (*i, j*) is transformed into a Weber contrast map,

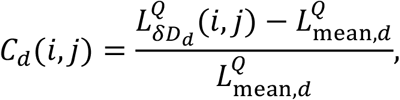

where 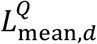 is the spatial mean of 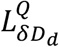 over the simulated retinal image. We then compute LSC at each pixel (*i, j*) within the RF centered on that pixel.

As in Karamanlis & Gollisch (2021) and Liu et al. (2022), we model this RF with a Gaussian kernel.

We sample this Gaussian on the retinal image grid, with *σ* expressed in micrometers on the retina and converted to *σ*_px_ through the retinal pixel size. Let *n*_*σ*_ be the half-size of the resulting kernel in pixels, and let *K*_*σ*_ = {(*k, l*) *∈* ℤ^2^: −*n*_*σ*_ ≤ *k, l* ≤ *n*_*σ*_} be the corresponding set of pixel offsets. Throughout the following derivation, *G*_*σ*_ denotes the sampled kernel normalized to unit sum:

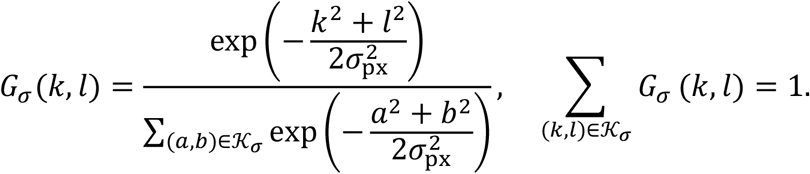

We use the weight *G*_*σ*_(*k, l*) for the sample *C*_*d*_(*i* + *k, j* + *l*). The local mean is

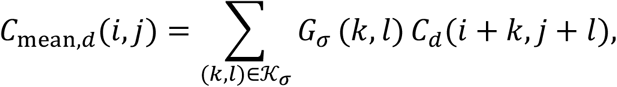

LSC is then defined as the weighted standard deviation of the Weber contrast values within the RF:

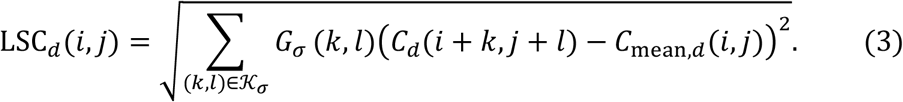

The local mean and local variance can also be written in terms of two local moments,

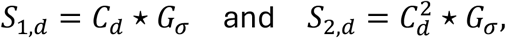

where *⋆* denotes cross-correlation. The LSC map is then

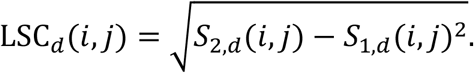

This expression is equivalent to Equation 3 and reduces the calculation to two cross-correlations per RF size and retinal defocus offset, which is computationally efficient with FFT.

The LSC-CARE relationship is evaluated for *σ ∈* [1,50] *µ*m. This sampling lets us identify which spatial integration ranges are most strongly associated with clinical efficacy, without assuming a specific retinal ganglion cell (RGC) type or eccentricity-dependent RF size.

For each lens, scene, RF size, retinal defocus offset, and optical configuration, this calculation gives a 2-D LSC map. We reduce each set of maps to its root-mean-square value across retinal positions and scenes. At offset index *d*, this reduction is

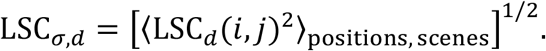

The LSC criterion used in the CARE analysis was then averaged across the *N*_*D*_ sampled retinal defocus offsets:

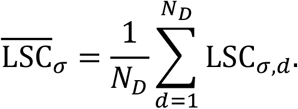

To assess the relationship with CARE, we divided the LSC value averaged over defocus for each MC spectacle lens by the corresponding value for the SVL under the same optical configuration and used this ratio in all regressions.

##### 4.5.1.1 Frequency-domain expression for local spatial contrast

The derivation of the frequency-domain LSC with numerical validation is provided in the Appendix (Section 5.4).

We performed the full image-based calculation for all indoor scenes in a reference condition with lowered gaze (20°), a 5 mm entrance pupil, and an on-axis source beam. We chose this configuration because the projected pupil lay entirely within the treatment area, allowing LSC to capture MC-layer effects across lenses.

We then used a frequency-domain approximation to evaluate additional imaging conditions, including straight-ahead gaze and peripheral eccentricities. Starting from the LSC definition (Equation 3) and averaging over the image dataset gives the following approximation for any unit-sum RF kernel *G*_*σ*_:

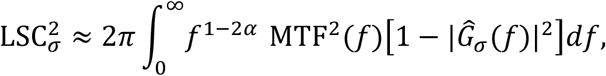

where MTF(*f*) is the radially-averaged MTF of the spectacle lens-eye system. For the continuous Gaussian approximation corresponding to this unit-sum RF kernel, |Ĝ_*σ*_(*f*)|^2^ = exp(−4*π*^2^*σ*^2^*f*^2^), which yields

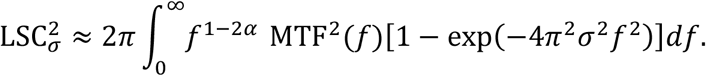

This relation is practically important: once the dataset spectrum and MTF are known, we can explore other datasets and source directions without computing convolutions for every image.

The correlations with 12-month CARE reported in Section 2.3 were evaluated across the sampled RF sizes and eccentricities. The regression shown in Figure 2, panel A, uses an RF size of 15 *µ*m and an eccentricity of 10° under lowered gaze (20°). For this regression, LSC was averaged across the sampled retinal defocus offsets.

### 4.6 Statistical Analysis

Clinical efficacy results for the same lens reported by different studies are first aggregated into a single CARE value. The resulting 13 lens-design values are then fitted by ordinary least squares, with each lens design contributing equally.

#### Aggregating clinical efficacy results across studies

Consider a lens ℓ for which 12-month CARE has been reported in *n*_ℓ_ studies, indexed by *k* = 1, …, *n*_ℓ_. Let *y*_ℓ*k*_ be the CARE value reported by study *k* and se_ℓ*k*_ its standard error. Each study contributes an inverse-variance weight 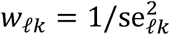. The aggregated CARE for lens ℓ is the inverse-variance-weighted mean

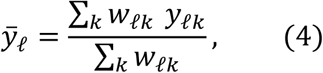

Each lens ℓ therefore contributes a single point 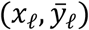, where *x*_ℓ_ is a metric extracted from the simulated retinal images, such as LSC_*σ*_. The standard error of 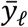, shown in Figure 11, is 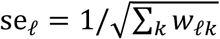.

#### Ordinary least-squares fit

We fit the per-lens points 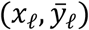 with the linear regression ŷ = *β*_0_ + *β*_1_*x*, giving every lens design equal influence:

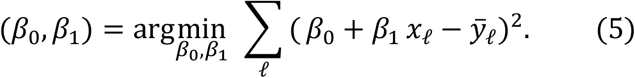

We perform the fit using statsmodels in Python. Fit quality is summarized with the coefficient of determination:

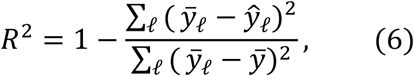

where ŷ_ℓ_ = *β*_0_ + *β*_1_*x*_ℓ_ are the fitted values and 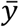 is the mean of the 13 aggregated CARE values.

#### Leave-one-out cross-validation

For the regression shown in Figure 2, panel A, we left out each lens design in turn, refitted the regression to the other 12 designs, and predicted CARE for the held-out design. We calculated the predicted residual sum of squares (PRESS) from these predictions. Leave-one-out cross-validated *R*^2^ was calculated as

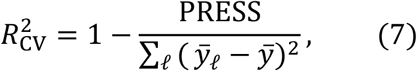

and leave-one-out cross-validated RMSE as

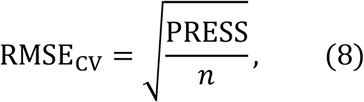

where *n* = 13. Because RMSE_CV_ is computed on CARE, it is expressed in millimeters.

#### Permutation test

We assessed the significance of the in-sample *R*^2^ by permutation. We reassigned the 13 aggregated CARE values at random among the designs. A single reassignment was applied across every sampled RF size, retinal eccentricity, and gaze direction, and *R*^2^ was recomputed at each. Within each reassignment we retained the highest *R*^2^ across this complete set of conditions, and we repeated the procedure 10 000 times. The reported p-value is the proportion of reassignments whose highest *R*^2^ equalled or exceeded the observed *R*^2^ for the regression shown in Figure 2, panel A, and is therefore adjusted for the full set of conditions examined. 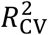 and RMSE_CV_ are calculated from the observed leave-one-out predictions and are not assigned permutation p-values.

## Competing interests

S.H., S.G., K.B., T.B. and S.M. are employees of EssilorLuxottica. O.M. has received financial support from EssilorLuxottica.

## 5 Appendix

### 5.1 Polychromatic point spread functions across lens designs

The figures below show the PSFs at the retina for the SVL and the 13 included MC spectacle lens designs. All calculations use a 5 mm entrance pupil, 3 D source proximity, 0.5 D accommodative lag, and lowered gaze (20°). The eye model was accommodated once through the on-axis SVL and reused for every lens and eccentricity.

**Figure 9.**
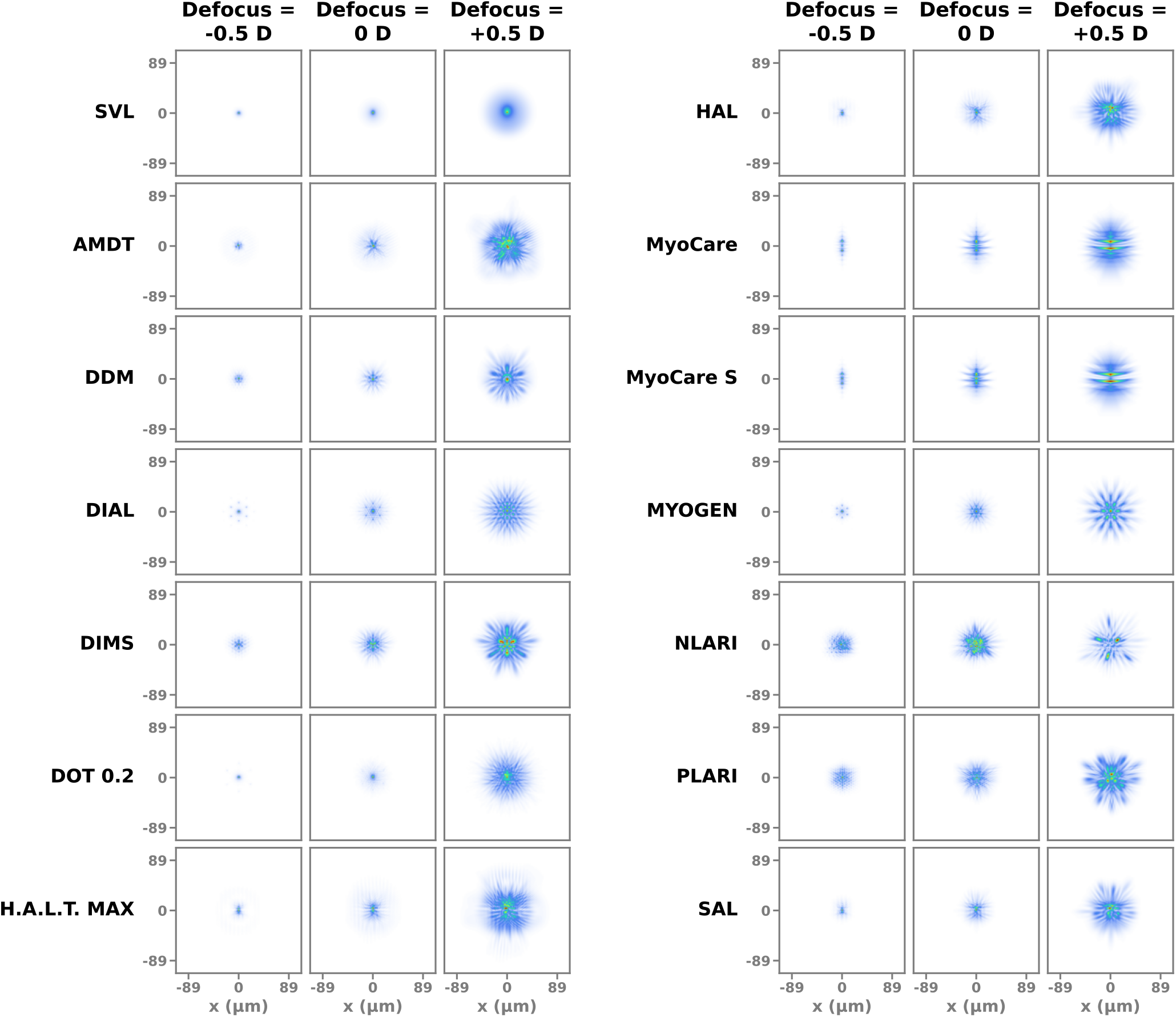
V(λ)-weighted polychromatic PSFs at the retina for the SVL and the 13 included MC spectacle lens designs, for an on-axis source imaged at the fovea under lowered gaze (20°).

**Figure 10.**
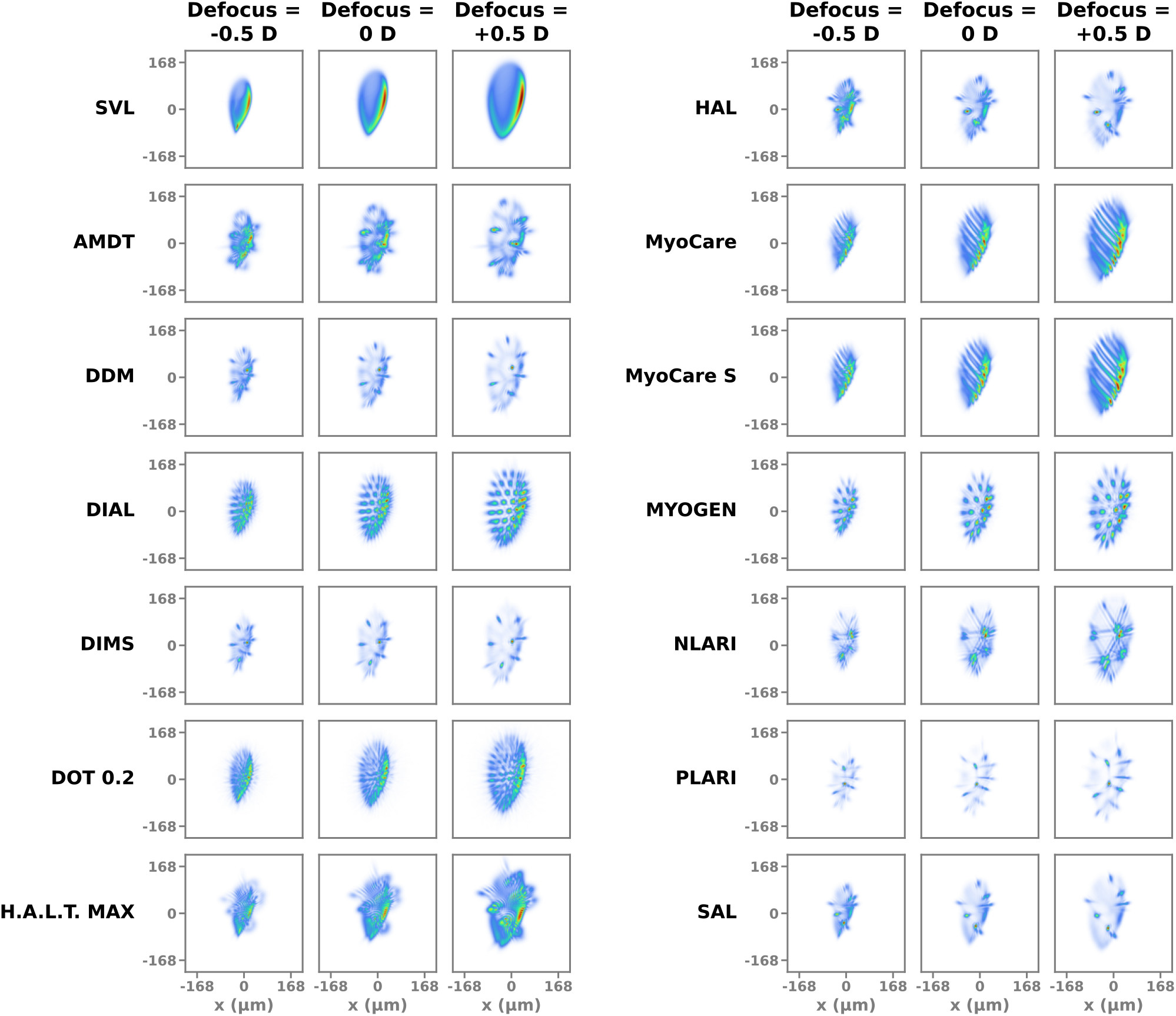
V(λ)-weighted polychromatic PSFs at the retina for the SVL and the 13 included MC spectacle lens designs, at 30^°^ retinal eccentricity under lowered gaze (20°).

### 5.2 Leave-one-out cross-validation

For each lens design in turn, the regression in Figure 2, panel A, was refitted to the other 12 designs and used to predict the CARE of the excluded design (Figure 11).

**Figure 11.**
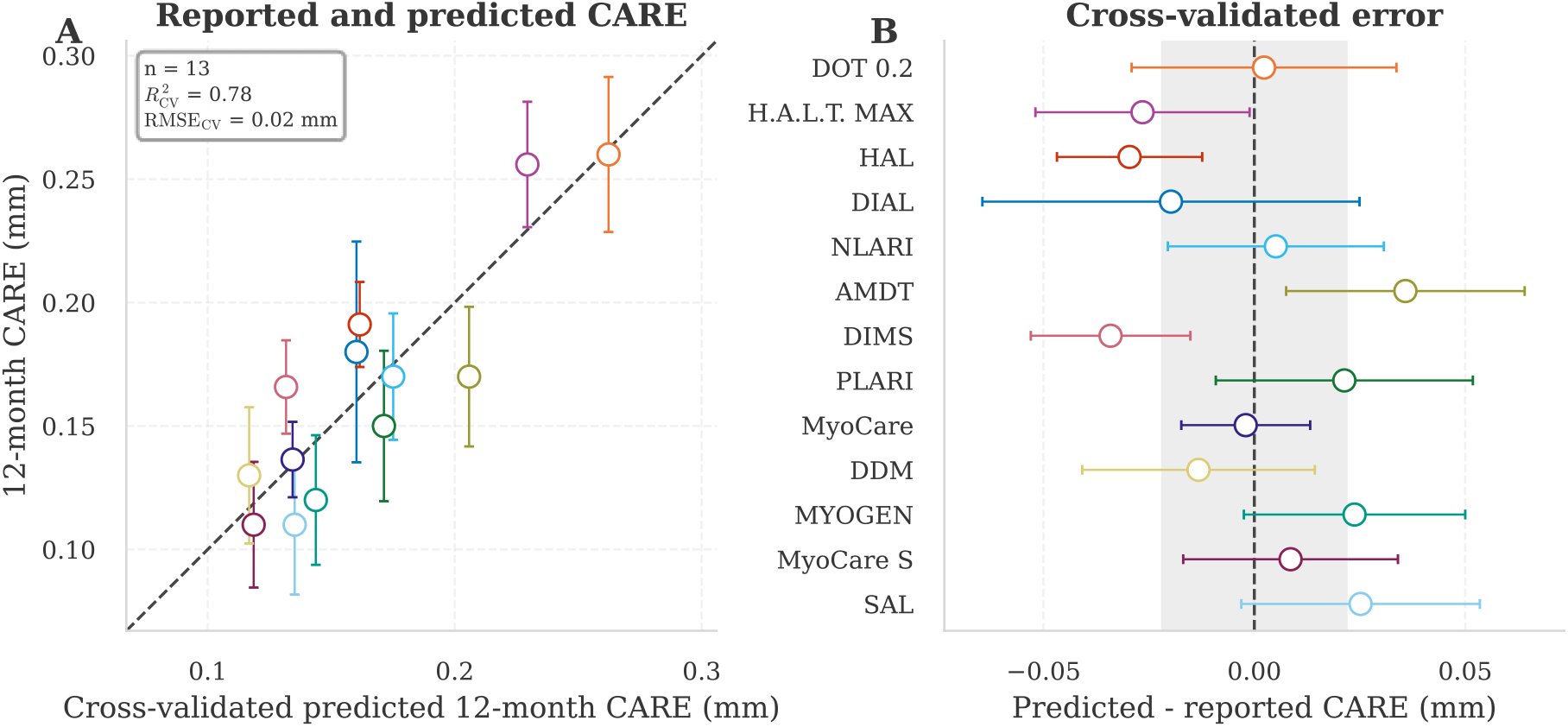
Leave-one-out cross-validation of the regression in Figure 2, panel A. CARE for each lens design was predicted from a fit to the other 12 designs. (A) Aggregated reported CARE against cross-validated predicted CARE. The dashed line shows equality. (B) Cross-validated error (predicted - reported CARE) for each design, sorted by reported CARE. The shaded band shows ±RMSE_CV_. Error bars show the standard error of the aggregated reported CARE (Section 4.6).

### 5.3 Entrance-pupil rays through the treatment area

Under straight-ahead gaze, ray tracing showed that the fraction of entrance-pupil rays passing through the treatment area increased from approximately zero at 10° eccentricity to 100% at 25° (Figure 12).

**Figure 12.**
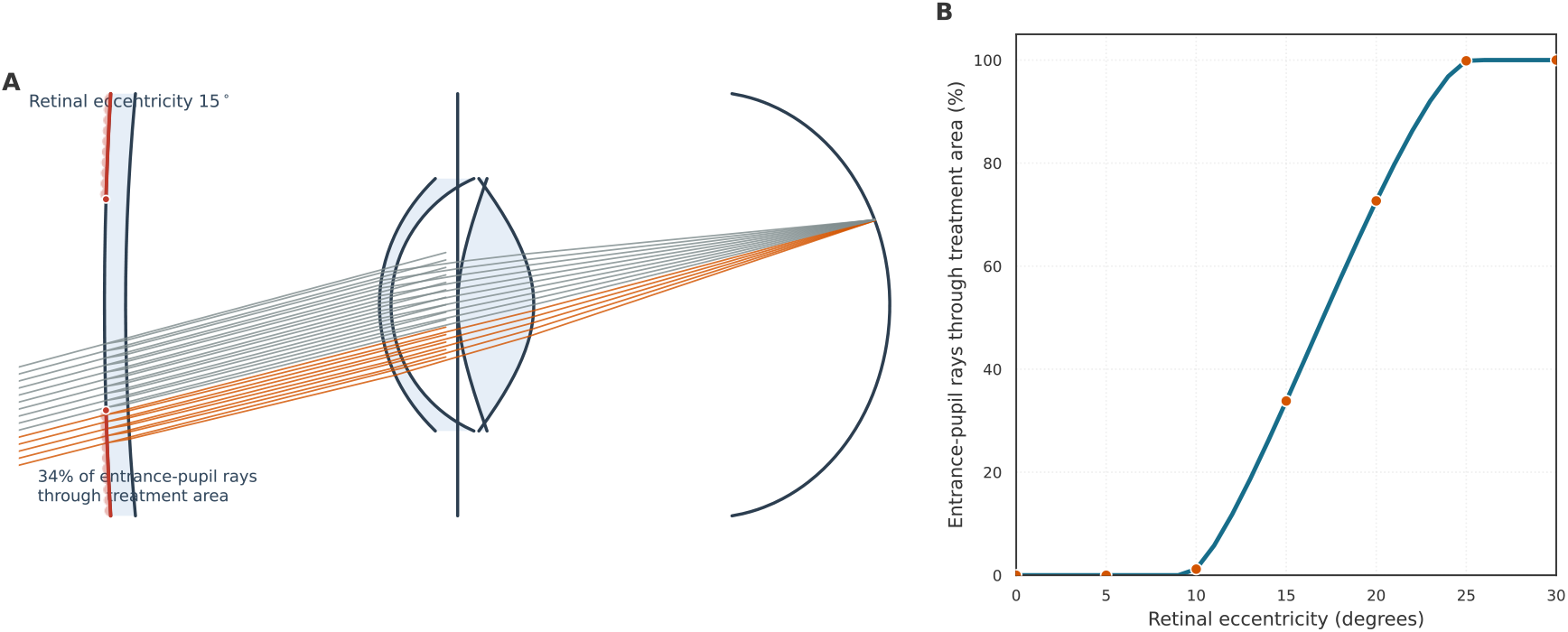
Fraction of traced rays passing through the treatment area as a function of retinal eccentricity. (A) Cross-section at an eccentricity of 15^°^. Gray and orange rays intersect the clear aperture and treatment area, respectively. The microlenses are shown schematically. (B) Fraction of a uniformly sampled entrance pupil intersecting the treatment area under straight-ahead gaze. Calculations used a 5 mm entrance-pupil diameter, 5 mm clear-aperture radius, 3 D source proximity, and 2.5 D accommodative response.

### 5.4 Local spatial contrast in the spatial-frequency domain

Computing LSC in the spatial domain over large image databases and multiple optical configurations can be computationally expensive, as each configuration requires image filtering followed by local contrast evaluation across the retinal image. To alleviate this cost, we derive a frequency-domain formulation of LSC that serves as a fast surrogate of the spatial metric.

#### 5.4.1 Theoretical framework

In this framework, image statistics are represented by the average power spectral density (PSD) of the indoor image dataset, while optical quality is characterized through the MTF. Under the assumption of wide-sense stationarity (WSS), the expected Weber-based LSC can be expressed as a weighted spectral-energy integral involving the image PSD and the RF transfer function. This formulation provides an efficient way to predict LSC variations across optical designs while preserving the dominant dependencies on image statistics and optical quality.

The derivation presented below should be considered a theoretical approximation rather than an exact description of real retinal images. Indoor images are neither strictly stationary nor ergodic, whereas the proposed formulation relies on a WSS representation of the Weber contrast field. Despite these simplifications, the resulting frequency-domain model captures the dominant second-order statistics relevant to LSC computation.

For clarity of notation, the following derivation is performed in the continuous domain. Let *C*(*r*) denote the continuous counterpart of the discrete Weber contrast map *C*(*i, j*), where *r* is a spatial position.

LSC is defined as

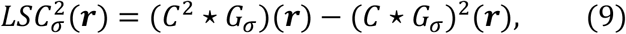

where *G*_*σ*_ denotes the normalized RF kernel and *⋆* the cross-correlation.

We further model *C*(*r*) as a WSS random process. Under this assumption, the mean of *C* is independent of spatial position,

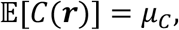

and the autocorrelation function

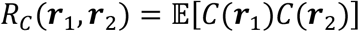

depends only on the spatial lag

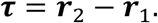

It can therefore be written as

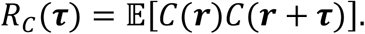

Our objective is to derive a frequency-domain expression of

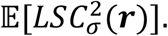

Using Equation 9,

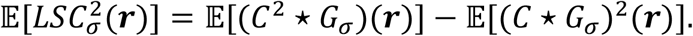

**First term**

Let us first evaluate

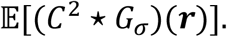

Since *G*_*σ*_ is deterministic,

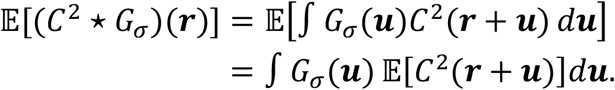

Since *C* is WSS,

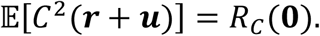

Therefore,

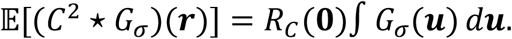

Using the normalization of *G*_*σ*_,

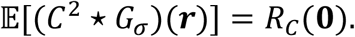

Applying the Wiener-Khinchin theorem,

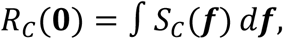

where *S*_*C*_(*f*) denotes the PSD of the Weber contrast field. Hence,

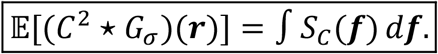

**Second term**

We now consider

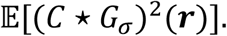

Let

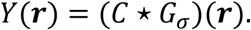

Since *C*(*r*) is WSS and *G*_*σ*_ is a deterministic linear shift-invariant filter, *Y*(*r*) is also WSS. The proof follows directly from the fact that the filtering operation preserves a constant mean and transforms the autocorrelation into a function of the spatial lag only.

To express this second-order moment in the frequency domain, we now derive the autocorrelation function of *Y*.

Since *Y* is WSS,

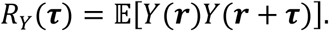

Substituting

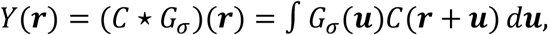

gives

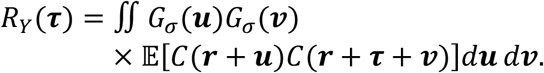

Using the WSS property of *C*,

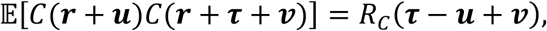

which yields

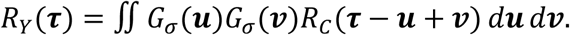

Recognizing the successive convolution operations and using the symmetry of *G*_*σ*_ yields

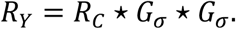

Applying the Wiener-Khinchin theorem to the WSS processes *C* and *Y*,

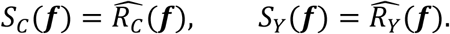

The convolution theorem gives

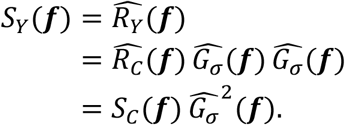

Since *G*_*σ*_ is a centered Gaussian kernel, 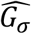 is real and positive. Therefore,

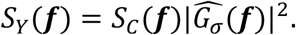

Applying the inverse Wiener-Khinchin relation at zero lag,

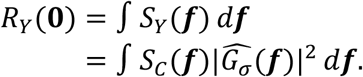

Since *Y* is WSS,

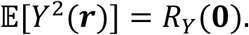

Therefore,

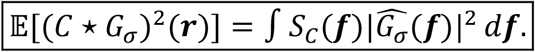

**Final frequency-domain formulation**

Combining the two previous results,

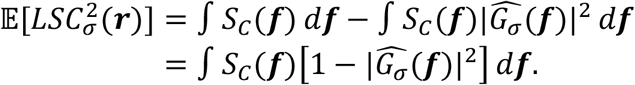

Assuming ergodicity, ensemble expectations can be approximated by spatial averages:

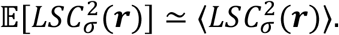

Therefore,

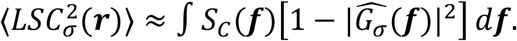

Finally,

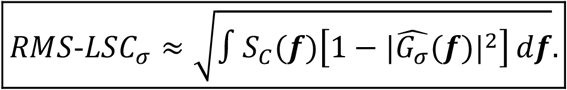

The quantity

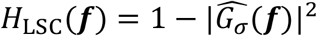

can be viewed as the effective high-pass weighting associated with LSC computation. Consequently, RMS-LSC squared corresponds to the spectral energy of the Weber contrast field weighted by this RF transfer function.

#### 5.4.2 Numerical validation of the WSS framework

The derivation presented in Section 5.4.1 assumes that the Weber contrast field is a WSS and ergodic random process. To assess the validity of the resulting frequency-domain formulation, we compare spatial- and frequency-domain LSC estimates on both synthetic WSS textures and real indoor images.

For each image and RF size *σ*, LSC was computed using both the original spatial-domain definition, *LSC*_spatial_, and the proposed frequency-domain formulation, *LSC*_freq_.

Synthetic WSS textures were first generated to satisfy the assumptions of the theoretical framework. The same analysis was then repeated on the indoor image dataset, which does not strictly satisfy the stationarity and ergodicity assumptions.

Figure 13 compares *LSC*_freq_ and *LSC*_spatial_ for all tested values of *σ*. Results from WSS textures and indoor images are shown in separate panels. For each dataset, a linear fit constrained to pass through the origin,

**Figure 13.**
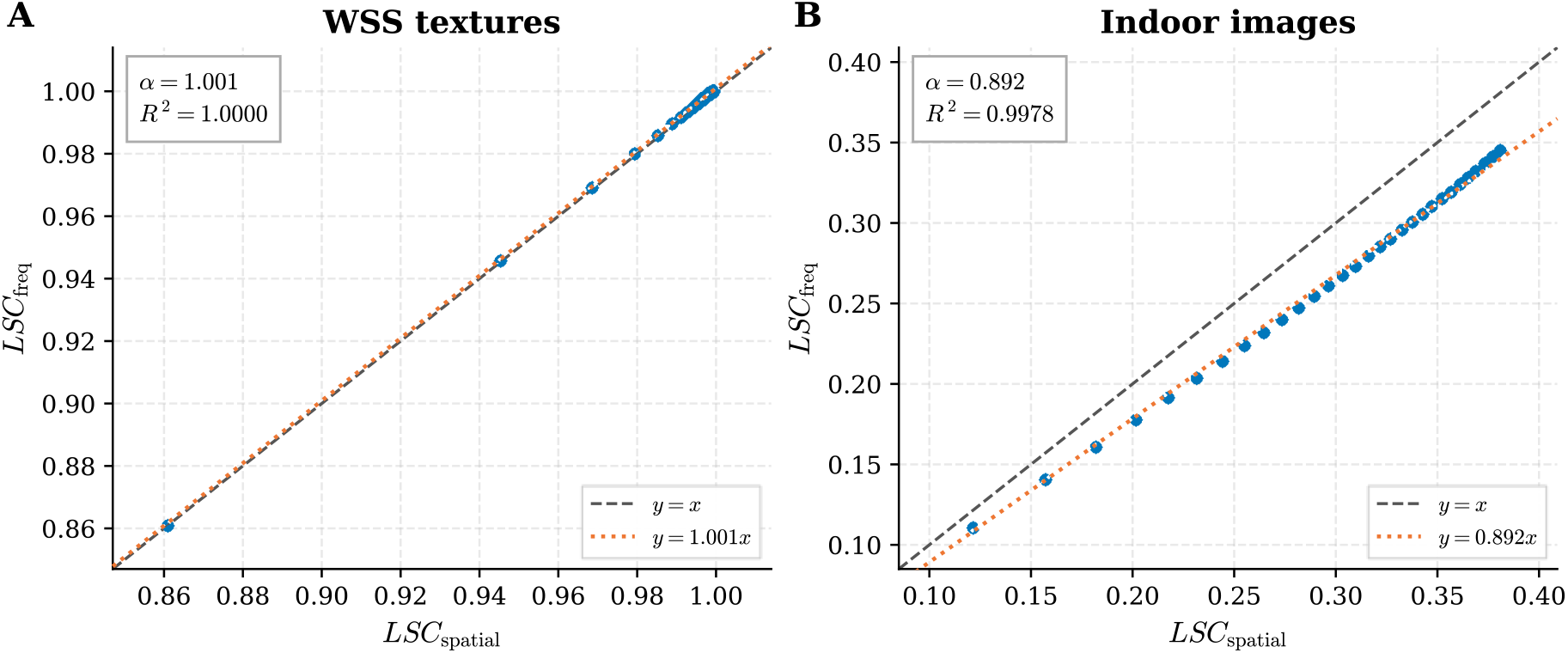
Comparison between spatial-domain and frequency-domain LSC estimates for synthetic WSS textures and indoor images. Each point corresponds to an image-σ pair. Linear fits constrained through the origin (LSC_freq_ = α LSC_spatial_) are shown together with the corresponding R^2^ values.

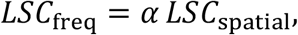

is also reported together with the corresponding coefficient of determination *R*^2^.

As expected, the WSS textures exhibit an almost perfect agreement between the spatial- and frequency-domain formulations, with a fitted slope close to unity and an *R*^2^ value approaching 1. This confirms the validity of the theoretical derivation under the assumptions introduced in Section 5.4.1.

Indoor images exhibit larger deviations due to departures from the WSS and ergodicity assumptions, as well as finite-image effects. Nevertheless, the relationship between both formulations remains strongly linear, as evidenced by the high coefficient of determination. The fitted slope of *α* = 0.892 indicates a systematic underestimation of the spatial-domain LSC by approximately 11%, while preserving a nearly proportional relationship between the two metrics. These results suggest that the frequency-domain formulation captures the dominant dependence of LSC on image statistics despite the simplifying assumptions introduced by the WSS framework.

Overall, the frequency-domain estimate is a reliable surrogate for spatial-domain LSC, both in the ideal WSS setting and with real indoor images.

We next extend the validated formulation to retinal image formation by introducing optical filtering and polychromatic image formation.

#### 5.4.3 Extension to optical image formation

The frequency-domain formulation derived in Section 5.4.1 can be extended to retinal image formation by incorporating the optical transfer function of the spectacle lens-eye system. The retinal image is obtained by convolving the object contrast field with the corresponding PSF, *PSF*_poly_. In the frequency domain, the retinal contrast PSD can therefore be expressed as

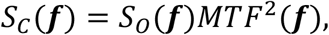

where *S*_*O*_(*f*) denotes the PSD of the object contrast field.

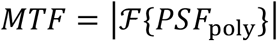

This MTF is associated with the retinal *PSF*_poly_.

Substituting this expression into the frequency-domain formulation of LSC yields

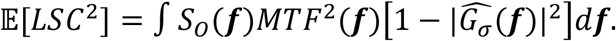

To further simplify the formulation, we approximate the two-dimensional polychromatic MTF by its radial average,

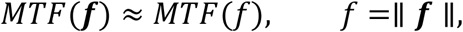

The RF transfer function is likewise assumed radially symmetric and, since *G*_*σ*_ is a Gaussian kernel, its squared modulus is given by

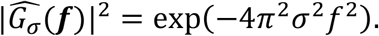

Likewise, the radially averaged spectral amplitude of natural images is often approximated by a power-law model,

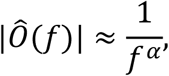

which implies

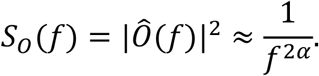

Under these assumptions, the frequency-domain formulation becomes radially symmetric. Expressing the integral in polar coordinates, *df* = *f df dθ*, and integrating over orientation yields

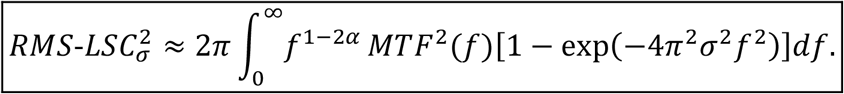

#### 5.4.4 Validation on realistic retinal image simulations

We performed a numerical comparison across multiple ophthalmic lens designs with RF sizes *σ* of approximately 15 and 30 *µ*m. For each optical configuration, the frequency-domain estimate was compared against the corresponding spatial-domain computation.

Figure 14 presents the relationship between the two metrics. Each point corresponds to a specific combination of indoor image, lens design, and RF size. Results for both RF sizes are displayed in the same figure.

**Figure 14.**
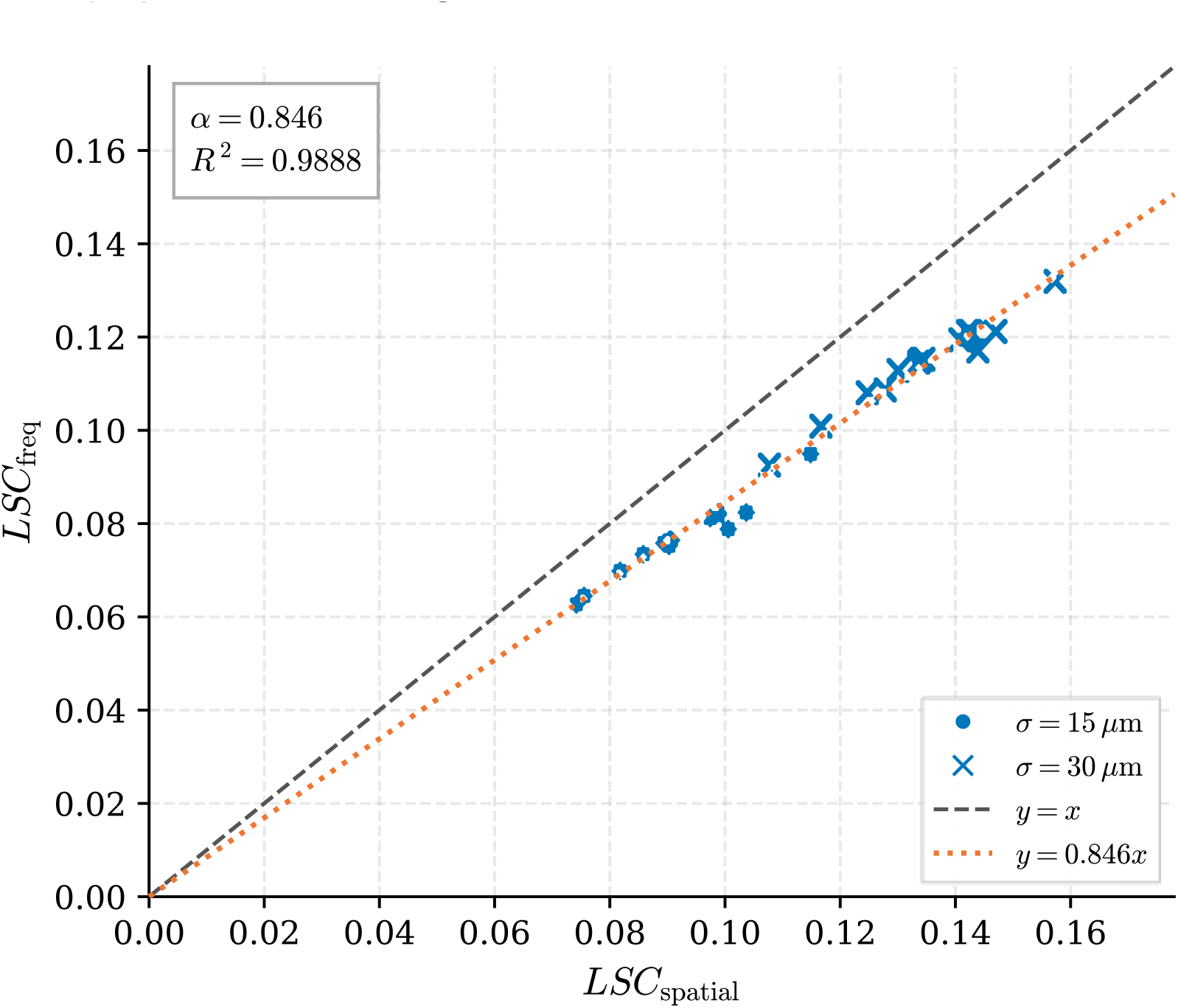
Spatial-domain versus frequency-domain LSC estimates for multiple ophthalmic lens designs evaluated with RF sizes σ of approximately 15 and 30 µm. The orange dotted line shows the best linear fit constrained through the origin (LSC_freq_ = 0.846 LSC_spatial_, R^2^ = 0.99). Despite a systematic scaling factor, the strong correlation validates the frequency-domain surrogate across realistic optical conditions.

We observe strong agreement between the spatial- and frequency-domain formulations across all investigated optical conditions. Despite the simplifying assumptions of the theoretical framework, the surrogate remains highly correlated with the spatial-domain LSC for both RF sizes. This indicates that the frequency-domain formulation preserves the dominant effects of image statistics and optical quality while providing a computationally efficient surrogate to evaluate ophthalmic lens designs.

